# Velamins: the first green-light emitting class of wild-type Ca^2+^-regulated photoproteins isolated from the ctenophore *Velamen parallelum*

**DOI:** 10.1101/2024.08.30.610560

**Authors:** Douglas M. M. Soares, Gabriela A. Galeazzo, Germán G. Sgro, Gabriela V. de Moraes, Leora Kronenberg, Emmanuella Borukh, Alvaro E. Migotto, David F. Gruber, John S. Sparks, Vincent A. Pieribone, Cassius V. Stevani, Anderson G. Oliveira

## Abstract

Ca^2+^-regulated photoproteins (CaPhs) consist of single-chain globular proteins to which coelenterazine, a widely distributed marine luminogenic substrate (the luciferin), binds along with molecular oxygen, producing a stable peroxide. Upon Ca^2+^ addition, CaPhs undergo conformational changes leading to the cyclization of the peroxide and the formation of a high-energy intermediate. Subsequently, its decomposition yields coelenteramide in an excited state and results in the emission of a flash of light. To date, all CaPhs reported produce blue light (λ_max_ 465-495 nm). Here, we report the cloning and functional characterization of a novel class of wild-type CaPhs capable of emitting green light: velamins, isolated from the bioluminescent ctenophore *Velamen parallelum*. Ten unique photoprotein-like sequences were recovered and grouped in three main clusters. Representative sequences were cloned, expressed, purified, and regenerated into the active His-tagged α-, β-, and γ-velamins. Upon injection of a calcium-containing buffer into the velamin, a flash of green light (λ_max_ 500-508 nm) was observed across pH values ranging from 7 to 9. Whilst α-velamin isoforms exhibited the highest light emission activity, β- and γ-velamins were found to be more thermostable at higher temperatures. Velamins are the only known wild-type Ca^2+^-regulated photoproteins that exhibit the longest wavelength in light emission, making them a promising model for studying spectral modulation. As a result, velamins hold potential for enhancing the sensitivity of signal detection in analytical systems, particularly when dealing with complex biological matrices.

## Introduction

Bioluminescence, the emission of visible “cold” light by living organisms, is a phenomenon widely distributed across kingdoms - from single-celled bacteria to vertebrates that exhibit complex and highly specialized light organs (1, 2). Interestingly, bioluminescent organisms such as annelids, cnidarians, cephalopods, gastropods, and ray-finned fishes, do not share a common mechanism of light production. In fact, it has been proposed that the bioluminescence trait has evolved independently at least 94 times across the tree of life (3). A typical bioluminescent reaction involves the oxidation of a substrate (luciferin), catalyzed by an enzyme (luciferase), generating electronically excited products that release energy in the form of photons when returning to their fundamental state (2). Alternatively, some luminous systems may include a luciferin–luciferase complex, known as photoprotein, that is more stable than its dissociated components (2, 4). Coelenterazine-photoproteins catalyze the addition of molecular oxygen into coelenterazine and stabilize the hydroperoxide formed within the protein (5, 6), requiring the addition of a regulating-element (usually a metal cation) to produce light (3, 7). Currently, more than 40 different types of bioluminescent systems have been elucidated, with at least half involving a photoprotein, including those sensitive to O_2_^•-^, H_2_O_2_, ATP, K^+^, and Ca^2+^ (4, 8, 9).

The majority of Ca^2+^-regulated photoproteins (CaPhs) are single-chain globular proteins of approximately 22 kDa to which coelenterazine, a widely distributed marine luminogenic substrate, binds within the protein’s internal cavity. Light emission is triggered by the binding of calcium ions, which induces conformational changes in the photoprotein. These changes initiate the decarboxylation of the 2-hydroperoxycoelenterazine moiety, resulting in the formation of coelenteramide in an excited state. The decay of excited coelenteramide to the ground state leads to light emission with a λ_max_ ranging from 465 to 495 nm (2, 5, 6, 10-13). To date, two classes of CaPhs from bioluminescent marine organisms have been the most extensively studied: one derived from hydromedusae (such as aequorin, mitrocomin, clytin and obelin) and the other from ctenophores (including mnemiopsin, berovin, bolinopsin and bathocyrovin). Despite sharing some common structural features, ctenophoran and hydromedusan photoproteins have low primary sequence identity and differ on certain intrinsic properties. For example, photoinactivation is present in ctenophores while hydromedusae photoproteins have a higher sensitivity towards calcium (14–16). In addition, the characterization of hydromedusan photoproteins is more extensive compared to ctenophoran CaPhs. CaPhs are one of the most versatile and sensitive tools to investigate Ca^2+^-regulated living processes for basic research and bioanalytical purposes. Cloning and expression of the genes encoding CaPhs from marine organisms such as *Aequorea victoria* (aequorin) and *Obelia longissima* (obelin), have opened new possibilities of applications, including highly-sensitive intracellular calcium indicators (13, 17). In imaging, CaPhs offer several advantages over calcium-fluorescent probes, since no external source of light is needed, avoiding photo-bleaching, phototoxicity, and autofluorescence issues (18, 19).

During the last two decades, major efforts have been taken to expand the collection of available photoprotein probes to improve their capacity as luminescent labels. In this report, we address these needs by isolating and characterizing a novel class of CaPhs: velamins, found in *Velamen parallelum*. This very active ctenophore species is characterized by a band-or wing-like gelatinous body that resembles a girdle, and it produces flashes of green light (λ_max_ 501 nm) when it comes into contact with something (20).

Specimens of *V. parallelum* were sampled along the southern Brazilian coast during a research expedition conducted aboard the MV Alucia vessel (Ocean X, USA) in the vicinity of the Alcatrazes Archipelago. DNA sequences encoding velamins exhibited high similarity to sequences of another bioluminescent ctenophore - *Mnemiopsis leidyi* - whose well characterized mechanism of light emission is also dependent on coelenterazine and Ca^2+^ (21). The new groups of photoproteins found in *V. parallelum* are the first wild-type CaPhs capable of emitting green light upon Ca^2+^ addition, holding promise for fundamental and applied studies on the molecular basis of spectral modulation in photoproteins. Furthermore, our results also suggest that some of these *V. parallelum* photoprotein isoforms could be exploited for their intrinsic thermostability and luminescence properties as an alternative bioanalytical reagent.

## Results and Discussion

### Ecology and molecular identification of *V. parallelum* specimens

The order Cestida includes only the family Cestidae, which comprises two monotypic genera: *Cestum* and *Velamen* (25), which are mostly oceanic and are usually found in the epipelagic (24) and mesopelagic zones (26, 27). They are found distributed globally across temperate, subtropical, and tropical seas.

The species *Cestum veneris* and *Velamen parallelum* are bioluminescent as are most pelagic ctenophore species. Nevertheless, *V. parallelum* emits greener bioluminescence (λ_max_ 501 nm) than *C. veneris* (λ_max_ 493 nm) (1, 20). They share similar general morphology and behavior. However, *V. parallelum* is smaller than *C. veneris* (which can reach more than 1 m), and is a more active species, undulating more vigorously and rapidly than *C. veneris* (23).

*V. parallelum* bioluminescence is usually manifested as flashes of light, always in response to a mechanical stimulus, *i*.*e*., when the animal touches or is touched by something. Considering their limited photosensitive capability, it is unlikely that *V. parallelum* bioluminescence is employed for intraspecific communication (20). Rather, it may have a function in avoiding predation: startling or blinding nocturnal predators (28), allowing the ctenophore to escape using its fast-undulating swimming mode (at a 90º angle to its cruising direction) to hide in the darkness.

During an expedition aboard the Research Vessel MV Alucia (Ocean X) close to the Alcatrazes Archipelago (São Paulo state, Southern Brazil), some bioluminescent ctenophores were observed swimming at a depth of around 500-600 m (Movie 1). These gelatinous animals were collected by using a manned submersible. A molecular approach involving DNA barcoding was used for a rapid identification of the specimens.

While the 18S rDNA gene, which encodes for the 18S ribosomal RNA, has been extensively employed to investigate broad relationships within ctenophores, this DNA barcode is highly conserved among different species and closely related genera, leading to poorly resolved trees (29–31). Indeed, analysis of the 18S rDNA gene sequenced from our specimens (GenBank accession OQ463859) revealed a significant degree of similarity with several sequences from *C. veneris* and *V. parallelum*, precluding their distinction (Fig. S1). Alternatively, the mitochondrial cytochrome-c-oxidase subunit 1 (COI) gene has been used in many phylogenetic studies, revealing a high diversity within many groups of Ctenophora which was previously not observed by using other DNA barcodes (30). In fact, by comparing the COI sequence from our specimens with those from other lobate ctenophores, it was possible to confirm the collected specimens as *V. parallelum* from a maximum likelihood analysis, as denoted by a high bootstrap support value in the nodes which segregate *C. veneris* from *V. parallelum* COI sequences (Fig. 1).

**Figure 1.**
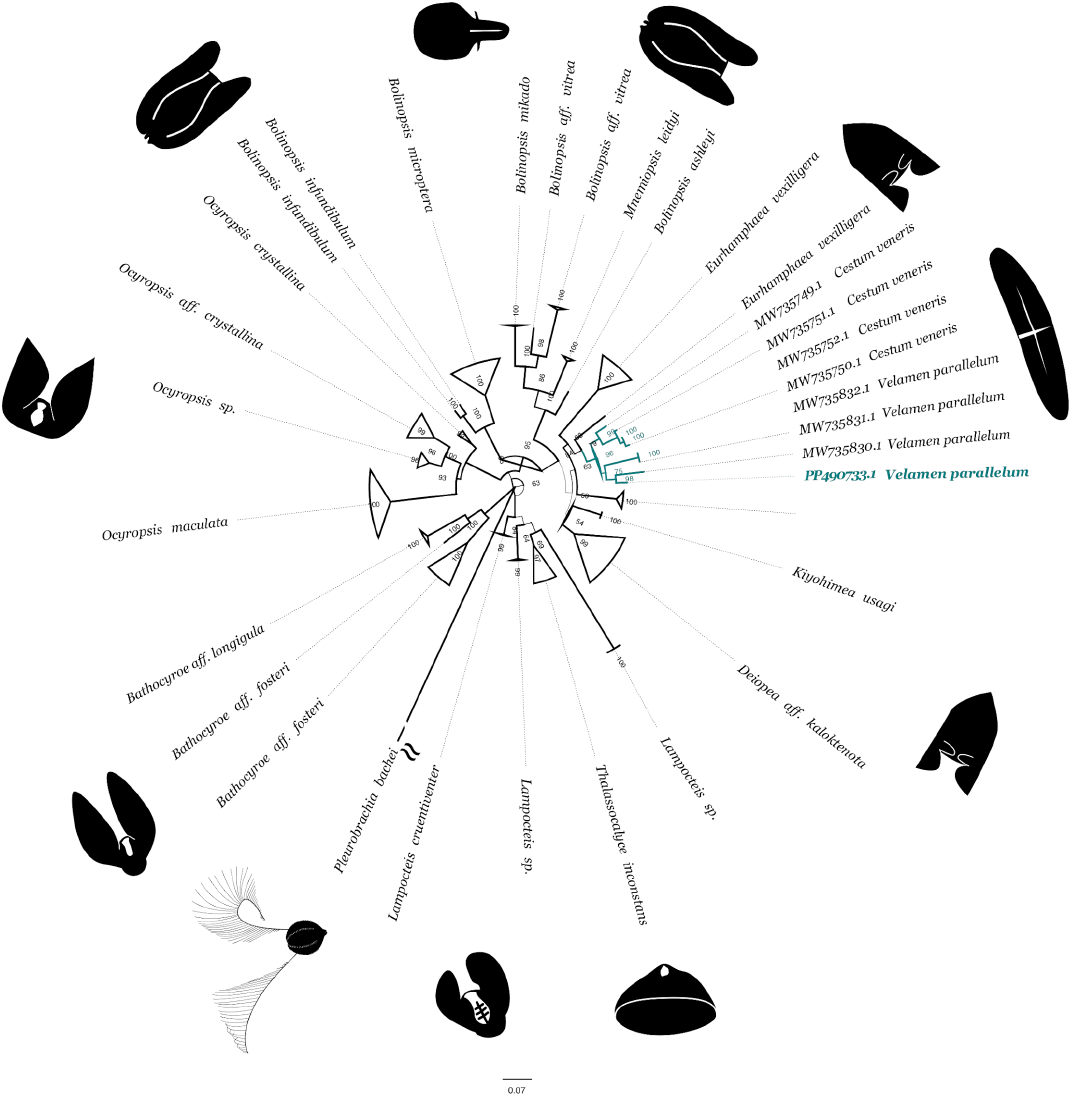
Maximum likelihood analysis of COI sequences from lobate ctenophores. Nodes corresponding to sequences from Cestidae specimens, including our *V. parallelum* specimen, are highlighted. Bootstrap support values are denoted in the branches. Silhouette images were obtained from PhyloPic (www.phylopic.org).

### Phylogenetic analysis of CaPhs

Analysis of 25 clones bearing photoprotein-like sequences of *V. parallelum* allowed for the recovery of nucleotide sequences coding for ten unique proteins - namely from VparPP1 to VparPP10, each one composed of 207 amino acid residues. A phylogenetic analysis was conducted using full-length amino acid sequence alignments of these proteins with those of other ctenophoran and hydromedusan CaPhs. Despite structural similarities, the analysis confirms that ctenophoran and hydrozoan sequences fall into two distinct clusters (Fig. 2). This is evidenced by a strongly supported monophyletic clade (99%) for hydromedusan photoproteins, which encompasses sequences from *Aequorea coerulescens, A. macrodactyla, A. parva, A. victoria, Clytia gregaria, C. hemisphaerica, Mitrocoma cellularia, Obelia geniculata*, and *O. longissima* (Fig. 2). These findings suggest that these proteins have undergone relatively rapid diversification since the divergence of ctenophores and cnidarians, as consistent with previous reports (21).

**Figure 2.**
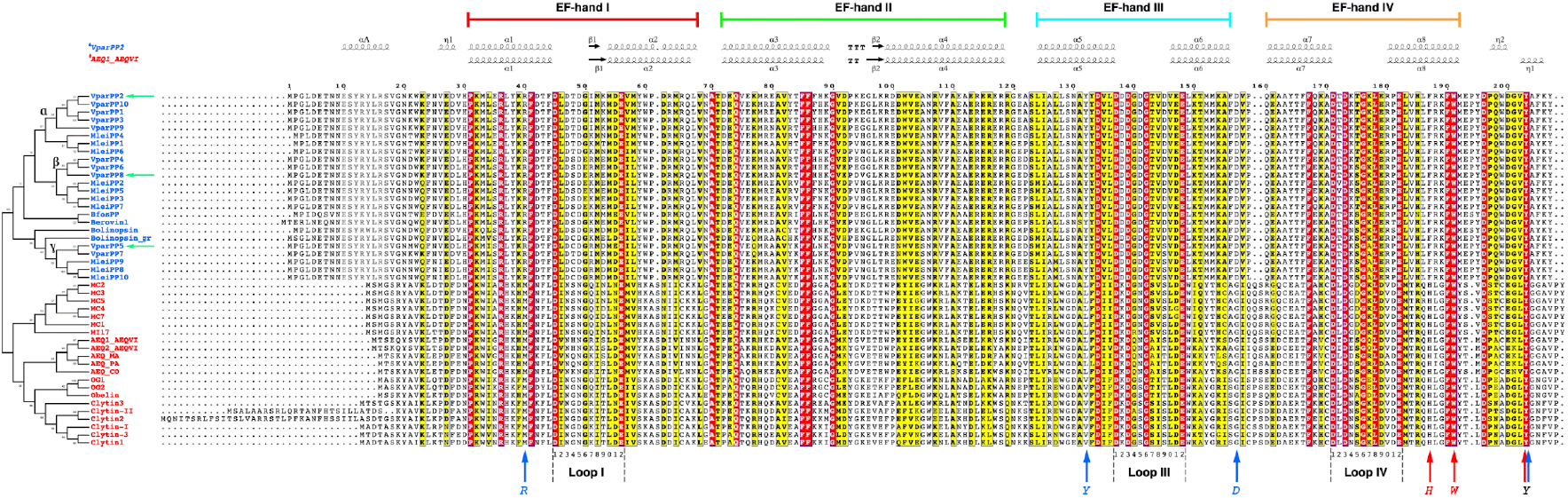
Maximum likelihood phylogeny of photoproteins from ctenophores (blue) and hydromedusae (red).

As observed in previous studies on ctenophores, photoprotein sequences of *V. parallelum* are also polyphyletic and grouped into three primary clusters along with mnemiopsin sequences, isolated from the ctenophore *M. leidyi* (21). The first one (α; 84% bootstrap support) includes the sequences VparPP1-3, 9, and 10, and forms a sister group to the mnemiopsin A (MleiPP1,4,6) clade. A second well-supported clade (β; 85%) contains the VparPP4, 6, and 8 sequences, closer to the mnemiopsin B sequences (MleiPP2,3,5,7). A third cluster (γ) comprises a branch including the VparPP5, 7, and mnemiopsin C (Mlei8-10) sequences.

### Sequences encoding CaPhs are conserved in Ctenophora

A multiple alignment analysis of the photoprotein sequences from the ctenophore *V. parallelum* revealed a high level of identity (from 86% to 99.5%) among them (Fig. 2 and Table S1), similar to those observed for mnemiopsin (87-100% among MleiPP1-MleiPP10) (21). In *M. leidyi*, it was demonstrated that these ten functional redundant mnemiopsin sequences are encoded in tandem by arrayed genes organized into two genomic clusters, which are likely maintained by strong purifying selection and concerted evolution (21). Persistent luminescence in *Mnemiopsis*, even under continuous stimulation, supports the hypothesis that maintaining multiple functional copies of photoprotein genes in the genome is an advantageous strategy. This ensures the constant synthesis of components required for light emission (21, 32, 33). However, due to the limited availability of genomic and transcriptomic data, it is not possible to infer the evolution and genomic structure of photoprotein genes among different taxa from Ctenophora.

Comparative analysis of the primary sequences (Fig. 2) also revealed a high degree of identity among ctenophoran photoproteins and illustrated that *V. parallelum* sequences are closer to mnemiopsin (85.5% to 96.1% identity) than sequences from other ctenophores, including those from *Beroe abssycola* and *Bolinopsis infundibulum* (84-90.3% of identity) (Table S1). In contrast, a lower degree of identity (8.7-27.1%) was observed among *V. parallelum* sequences and hydromedusan photoproteins, including aequorin, clytin, mitrocomin, and obelin (Table S1). These findings are consistent with differential bioluminescence properties between ctenophoran and hydromedusan photoproteins, such as photoinactivation in ctenophoran photoproteins and a higher sensitivity to calcium reported for hydromedusan photoproteins.

The predicted molecular weight for the *V. parallelum* photoprotein sequences ranged from 24.49 to 24.74 kDa, with isoelectric points ranging from pH 4.31 to 4.71 (Table S2). As expected due to the high level of genetic similarity, these values are close to those reported for mnemiopsin sequences - with molecular weight ranging from 24.56 to 24.76 kDa and isoelectric points at pH from 4.57 to 4.82 (21).

In order to further characterize *V. parallelum* photoproteins (VparPPs), we cloned *E. coli*-optimized versions of VparPP2, VparPP8, and VparPP5 into pET28a vectors to obtain the recombinant α-, β-, and γ-apovelamins, respectively.

### Functional analysis of velamins

The highest light emission intensity was observed for α-velamin, followed by β- and γ-velamin (Fig. 3A). After the initial characterization, optimal conditions for light emission were determined by monitoring light emission activity using different buffers. The results showed that velamins are most active in Tris-based buffers with a pH range of 8.5-9.0 (Fig. 3A), which is similar to the optimal pH values reported for other ctenophoran photoproteins, such as recombinant bathocyrovin (pH 8.5) from *B. fosteri* (16) and native and recombinant mnemiopsins (pH 9.0) (21, 34).

**Figure 3.**
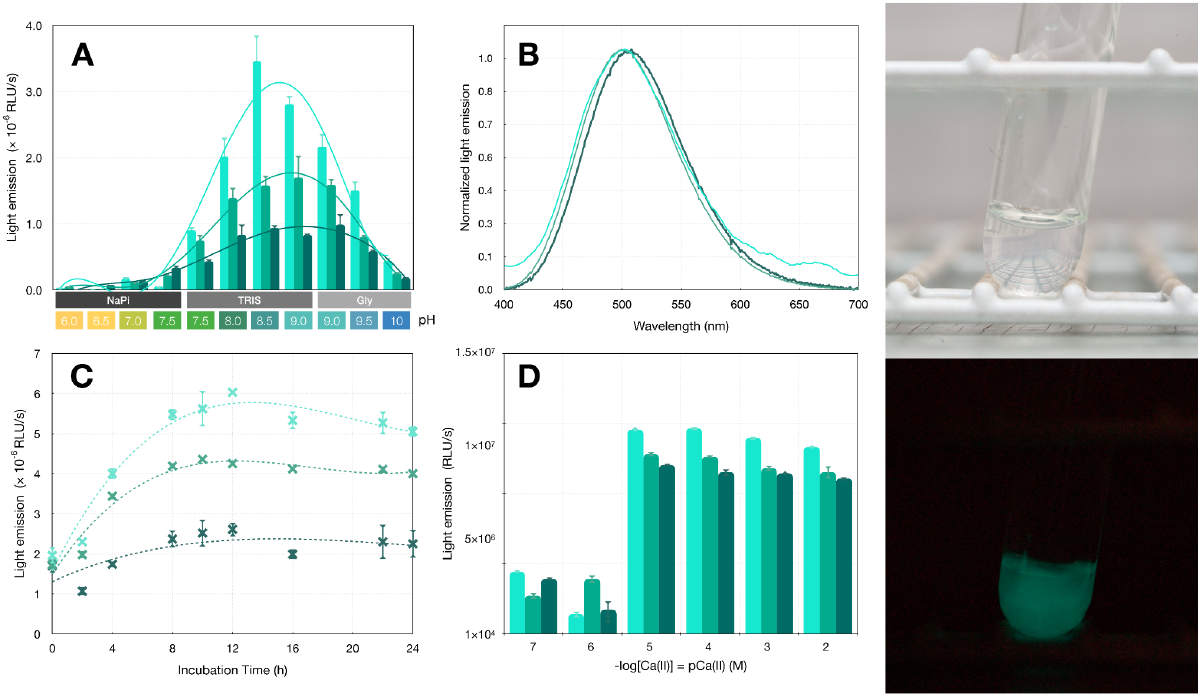
Characterization of recombinant α-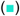, β-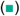, and γ-velamin 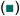. (**A**) Chemiluminescence assays in different buffers with pH ranging from 6 to 10; (**B**) spectra of light emission; (**C**) the influence of different incubation times on chemiluminescence intensity; (**D**) response of velamins to different concentrations of calcium ions. *Inset*: magnification of pCa(II) 7 and 6. Test tubes: a flash of green light observed right after the addition of Ca^2+^-supplemented buffer to the velamins.

A flash of green light was observed immediately after the addition of Tris buffer supplemented with calcium chloride into the test tubes containing velamins (Fig. 3 and Movie 2). The light emission wavelength observed for γ-velamin (λ_max_ 508nm) was slightly red shifted compared to the α- and β-velamins (λ_max_ 500nm) (Fig. 3B). A similar result was reported for the mnemiopsin C MleiPP9 (λ_max_ of 496 nm at pH 8), equivalent to γ-velamin, in comparison to the mnemiopsin representatives of groups A and B (MleiPP6 and Mlei_PP3, respectively, with λ_max_ of 490 nm) (21). No significant changes in the spectra of light emission were observed in chemiluminescence tests conducted with velamins in a pH range from 7 to 9 (Fig. S4). Similarly, the blue light emitted by mnemiopsins displayed only minor variations of 1 nm across a pH range of 8 to 10, as previously reported (21).

Ctenophoran photoproteins, similar to those of hydromedusae, can be reconstituted from apoproteins through the incubation with coelenterazine under calcium-free conditions in the presence of oxygen. However, there is a distinction in the conversion process: ctenophoran apophotoproteins require alkaline pH solutions and high ionic strength, whereas hydromedusan apophotoproteins can be efficiently transformed into active proteins at neutral pH and low ionic strength (10). We conducted an evaluation of various incubation times during the regeneration process of velamins in the presence of coelenterazine. Our findings revealed that apovelamins can undergo complete regeneration into their active forms within a period of 8 hours (Fig. 3C). The time taken for regeneration is half of what is needed for mnemiopsins to complete the process (21).

Considering the extensive utilization of hydromedusan CaPhs as tools for intracellular calcium measurements, we also investigated the response of velamins to different calcium concentrations (Fig. 3D). The results demonstrated that velamins had lower calcium sensitivity than aequorin and other hydromedusan CaPhs, which is an intrinsic feature of ctenophoran CaPhs. As previously discussed (10, 35), a proper comparison among ctenophoran CaPhs is limited due to the existing calcium concentration–effect curves for mnemiopsins, which do not adequately allow for an objective assessment of the changes in light intensities at low [Ca^2+^]. Therefore, additional studies are necessary to address this gap in knowledge.

Following velamin characterization, we conducted an evaluation of the thermal stability of these proteins. Considering the superior performance of α-velamin in terms of light emission, we investigated its stability at 37ºC, which would be vital for further studies aiming the application of this CaPh for *in vivo* bioluminescent assays. A progressive decrease in light emission was observed from the incubation of the regenerated α-velamin at 37ºC, being almost depleted in about 10 minutes (Fig. S5A).

Additionally, we also investigated the thermostability of the apovelamins by incubating them at temperatures ranging from 25 to 75°C for different time periods. Following the incubation, the samples were then placed on ice for 30 minutes before being regenerated into their active velamin forms for further analysis through light emission assays. The objective of this experiment was to examine the residual activity of the photoproteins, demonstrating the degree to which they maintain their functionality following structural modifications resulting from heating in their apo form. The findings from thermal inactivation tests indicated no significant alterations in the activity of β- and γ-velamins after incubation at 25 and 37ºC for up to 60 minutes (Fig. 4). A decrease of approximately 20% in the activity of α-velamin was observed under the same experimental conditions (Fig. 4). Considering the initial normalized chemiluminescence, the performance of the β- and γ-velamins suggests that they are promising to be investigated in further studies of thermostability.

**Figure 4.**
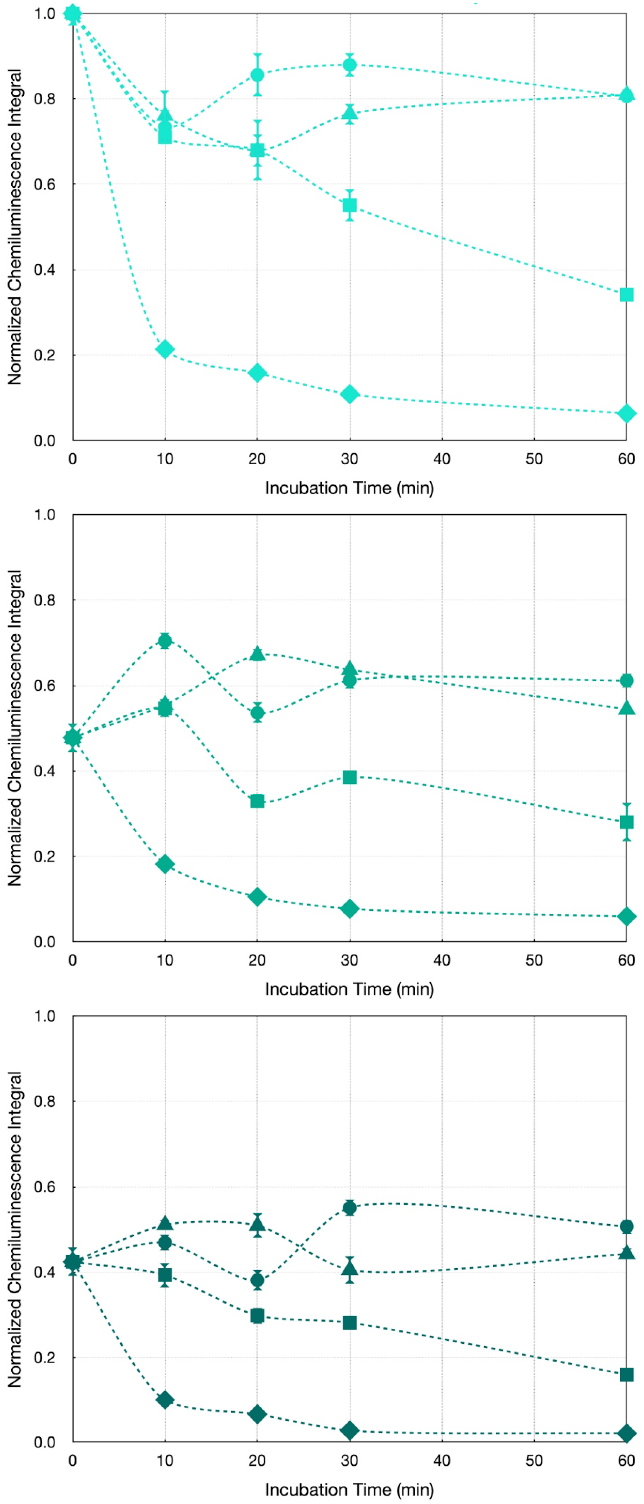
Normalized chemiluminescence of α-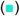, β-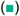, and γ-velamin 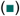 in assays of thermal inactivation after the incubation of the apophotoproteins in different temperatures (• 25ºC, ▲ 37ºC, ▪ 50ºC and ◆ 75ºC) and time periods. Average and standard deviation are calculated from the estimation of integral values obtained for technical replicates.

In a recent study, similar findings were reported when comparing the thermostability of apomnemiopsin (ctenophore) to apoaequorin (hydromedusa) (36). Following a 60-minute incubation at 40ºC, apomnemiopsin retained 71.4% of its initial activity. Remarkably, even after being subjected to a 30-minute incubation at 80ºC, a considerable level of activity was still maintained. The authors also demonstrated that while apomnemiopsin and apoaequorin have similar tertiary structures, apomnemiopsin exhibits higher stability at elevated temperatures. This increased stability can be primarily attributed to a favorable enthalpy contribution. Furthermore, the presence of favorable residues and the number of interactions, which are sequence-based parameters, may also play a significant role in determining the thermal stability of CaPhs (36).

Moreover, photoinactivation assays made with α-velamin exposed to ambient light revealed a decline in emitted light over time (Fig. S5B), consistent with findings reported for other ctenophoran CaPhs (16, 21, 37). Tests of photoinactivation were also conducted by exposing the velamin to UV light, which resulted in complete photoinactivation of the light emission after 5 minutes (data not shown).

### Structural comparisons among CaPhs

Despite their low primary sequence similarity, ctenophoran and hydromedusan photoproteins exhibit a high degree of structural similarity. Remarkably, ctenophoran photoproteins possess an additional N-terminus α-helix (herein called αA) that includes a highly conserved motif, ESY(R/K)(Y/W)LRS (positions 10-17 in VparPP2; Fig. 2), which is absent in hydromedusan photoproteins (Fig. S6 and Movie 3). In order to assess the relevance of this secondary structure for the function of ctenophoran CaPhs, we created a shorter α-velamin version lacking the ESYRYLRS motif. Despite the successful expression and purification of the respective apovelamin, no light was detected after calcium addition to this velamin mutant (data not shown). This finding demonstrates this N-terminus α-helix is crucial for the function of ctenophoran CaPhs, although its mechanistic role remains unclear. Additionally, the identification of this strictly conserved motif contributes as an additional feature to identify new ctenophoran photoproteins as well as distinguish them from hydromedusan CaPhs.

Ctenophoran and hydromedusan CaPhs also share four highly conserved EF-hand domains (I to IV), each made of a helix-loop-helix (HLH) motif, summing at least 8 α helices (Fig. 2). Random mutagenesis and functional screening of aequorin mutants suggest that single EF-hands are capable of triggering luminescence at a slow rate, with higher calcium affinity to the C-terminal EF-hand pair compared to the N-terminal EF-hand (15).

In a closer look, all calcium-binding domains at EF-hands I, III, and IV are organized in a 12-amino acid central loop, corresponding to residues 45-56 (EF-hand I), 137-148 (EF-hand III) and 171-182 (EF-hand IV) of α-velamin (VparPP2) (Fig. 2, highlighted as loops I, III, and IV). Calcium-binding residues at loop positions 1 and 12 are strictly conserved in all EF-hand domains, occupied by Asp and Glu residues, respectively. Residues at loop positions 3, 5, 7, and 9 are also capable of binding calcium. The 3^rd^ position is typically occupied by Asp, except for an Asn substitution occurring solely in the EF-hand I of hydromedusan photoproteins (Fig. 2). An Asp residue is also commonly observed at the 5^th^ loop position in ctenophoran EF-hands I and III, although it is replaced by Ser/Thr residues in the EF-hand IV (Fig. 2). On the other hand, amino acids at the loop positions 7^th^ and 9^th^ are hypervariable; except for the 7^th^ position of EF-hand III in ctenophores, occupied by a conserved Thr, and the 9^th^ position of EF-hand IV, where a conserved Glu occurring in ctenophoran photoproteins is replaced with an Asp in hydrozoans (Fig. 2).

Furthermore, the 6^th^ position of the calcium-binding loops (50, 142, and 176 in VparPP2) is occupied by a conserved Gly residue in all CaPhs, except for its replacement by a Glu at the EF-hand I of mnemiopsin B (MleiPP2, MleiPP3, MleiPP5, MleiPP7) and β-velamin (VparPP4, VparPP6, VparPP8) sequences (Fig. 2). The functional relevance of this conserved residue was investigated in mnemiopsin B variants Gly142Glu (loop III), Gly176Glu (loop IV), and a double mutant (38). Overall, it was demonstrated that changes at loops III and IV affect the decay rate and the intensity of light emission, respectively. Therefore, it suggests that the evolutionary selection of Gly at the 6^th^ position of those loops is essential for achieving an optimal balance between these features in ctenophoran CaPhs (38). Similar findings were reported from Gly36Arg (loop I) and Gly129Arg (loop III) aequorin mutants, whose amino acid replacements at the 6^th^ position of loops I and III resulted in losing all and half of aequorin activity, respectively (39).

The contribution of other residues on the differential bioluminescence properties between ctenophoran and hydrozoan photoproteins has also been investigated. For instance, the replacement of the 7^th^ residue at the first α-helix in EF-hand II of ctenophoran CaPhs for other one commonly found in hydrozoan CaPhs resulted in the complete loss of luminescence in the Met77His mnemiopsin (MleiPP1) mutant, demonstrating the functional relevance of this residue in ctenophoran photoproteins (40). Similarly, Trp101Phe, Trp101Tyr (4^th^ residue of the second α-helix in EF-hand II), and Met151Tyr (7^th^ residue of the second α-helix in EF-hand III) mnemiopsin (MleiPP1) mutants showed significant alterations in luminescence activity, decay rate, and calcium sensitivity (40).

Site-directed mutagenesis studies from hydromedusan photoproteins have offered new insights into understanding the functional roles of specific residues at the corresponding positions in ctenophoran photoproteins. For example, Cys158Ala obelin mutant, with a substitution of a conserved Cys for the ctenophoran equivalent Ala (residue right before loop IV; Ala170 in VparPP2; Fig. 2), as well as the Cys158Ser mutant, showed altered chemiluminescence properties and slowed photoprotein complex formation in the series: wild type>Cys158Ser>Cys158Ala, consistent with the nucleophilicity of their side chain (SH>OH>CH_3_) functional groups (41).

The contribution of α-helices to the stabilization of hydromedusan photoproteins has also been proposed based on the crystal structures of aequorin and obelin. Modifications at the conserved residue Arg21 (Arg36 in VparPP2), located at the first α-helix of EF-hand I and with which the C-terminal Pro195 (208^th^ residue in the alignment of Fig. 2) interacts via a hydrogen-bond network, have a detrimental effect on light emission and stability of the photoproteins (42). Unlike the proline residue, this arginine is highly conserved among all CaPhs. However, the lack of a known molecular structure for the C-terminal end of MleiPP1 and Berovin1, the only two photoproteins sequenced so far (Table S3), plus differences in the sequences, make it difficult to infer a similar interaction feature and effect.

Predicted structures of ctenophoran photoproteins suggest that their substrate-binding cavity contains fewer aromatic residues compared to cnidarians, with loops I and II being more negatively charged, which could result in different calcium sensitivity between the photoproteins of these two phyla (34). Hydromedusan CaPhs structure modeling proposes that the conserved triad His175-Trp179-Tyr190 in obelin and His176-Trp180-Tyr191 in aequorin interact with the peroxide and carbonyl groups of the 2-hydroperoxycoelenterazine through hydrogen bonds, as supported by studies using substitution mutants with different hydrogen bond donor-acceptor capacity and charged properties (43). The involvement of these residues in triggering chemiluminescence has also been demonstrated from studies which investigated the proton transfer from Tyr190 to His175 in obelin (44); and for His176Ala, His176Phe, His176Tyr aequorin mutants, which resulted in decreased luminescence activity (45).

In ctenophores, this triad is relatively conserved, except for the substitution of His at the first position with Phe, giving the triad Phe187-Trp191-Tyr203 in VparPP2. Interestingly, AlphaFold protein structure predictions suggested the ctenophoran triad Phe-Trp-Tyr has a similar spatial location/orientation to the His-Trp-Tyr triad found in hydromedusan CaPhs (Fig. 2, Fig. S7 and Movie 4). Unfortunately, the only two structures resolved for the ctenophoran CaPhs berovin and mnemiopsin (MleiPP1), (both apo-versions with a “built-in” peroxy adduct of coelenterazine) do not clearly show the final C-terminus residues, including the berovin Tyr204.

Additionally, a study combining structural modeling and the impact of substitutions on the chemiluminescent properties of berovin speculated that coelenterazine is bound as a 2-peroxy anion adduct within the internal cavity of ctenophoran photoproteins (37). As such, it is stabilized by a Coulomb interaction with a positively charged guanidinium group of Arg41 and the Tyr204. Then, calcium-induced conformational changes at berovin disrupt this interaction and result in light emission (37).

In addition to Arg41 and Tyr204 residues, quantum mechanics/molecular mechanics (QM/MM) simulations for berovin -and Cd^2+^-loaded apomnemiopsin (MleiPP1) suggest that these residues interact with Asp158 to form a conserved triad in ctenophoran photoproteins (46). This triad creates a hydrogen-bonded network that facilitates the transfer of a proton from the chromophore coelenterazine’s 2-hydroperoxy group to bulk solvent (46, 47). Interestingly, the coelenteramide binding cavity of mnemiopsin exhibits hydrogen bond networks that mediate the interaction of a water molecule with the amide nitrogen atom of coelenteramide, and between coelenteramide and an extra residue (Tyr131) via another water molecule (47). This supports the hypothesis that a coelenteramide-bound water molecule catalyzes the 2-hydroperoxycoelenterazine decarboxylation reaction by the protonation of a dioxetanone anion, triggering light emission (47). Both the triad Arg-Asp-Tyr and the extra tyrosine residue are found in all velamins described here (Arg40-Asp157-Tyr203 and Tyr132 in VparPP2) (Fig. 2, Fig. S8 and Movie 5).

Residues involved with the red-shifted spectra of γ-velamins and mnemiopsins C remain unclear. Three amino acid residues exclusive to these photoproteins were identified when compared to other ctenophoran photoproteins: threonine at position 98 that is replaced by lysine or leucine in other ctenophores; serine at position 145 that is changed for aspartic acid; and alanine at position 177 replaced by lysine (Fig 2). All of these residues are located between the two helices which compose the EF-hands II (Thr98 in VparPP5), III (Ser145 in VparPP5), and IV (Ala177 in VparPP5). Furthermore, residues 145 and 177 correspond, respectively, to the 9^th^ and 7^th^ positions of the calcium-binding loops (Fig 2). Experimental data obtained from the Val144Ile mnemiopsin-2 mutant also suggest the relevance of the conserved 10^th^ residue of the calcium-binding loop at the EF-hand III (Val146 in VparPP2, Fig. 2) regarding the function of ctenophoran CaPhs, whose substitution resulted in a lower initial intensity, slower decay rate, and a red shift of 25 nm in light emission (48).

Some hydromedusan CaPh mutants have suggested the involvement of the residue Tyr82 (aequorin)/Phe88 (obelin), equivalent to the Trp102 in VparPP2 (Fig. 2), on the differential light emission spectra of photoproteins. Whilst the Tyr82Phe aequorin mutant had its chemiluminescence redshifted from 469 nm (wild-type) to 501 nm, obelin mutants Phe88Tyr (λ_max_ 453 nm), Phe88Trp (λ_max_ 477 nm) and Phe88His (λ_max_ 459 nm) exhibited shorter light emission wavelengths in comparison to the wild-type (λ_max_ 482 nm) (49). These alterations in the wavelength could be explained by the interaction between the aequorin Tyr82 and the coelenteramide via H-bond, absent in obelin (49). Crystal structures of two conformational states of the Phe88Tyr obelin revealed that this substitution does not affect the overall protein structure compared to the wild-type version and corroborate the previous hypothesis that H-bond patterns near the oxygen of the 6-(p-hydroxyphenyl) substituent of coelenterazine are the basis for spectral differences among photoproteins (50).

Further studies are required to explore the significant differences between ctenophoran and hydromedusan photoproteins, in order to investigate the specific residues or secondary structures that contribute to their intrinsic bioluminescence properties. Ctenophoran photoproteins may present some advantages over hydromedusan photoproteins, as they do not require reducing agents for activation and are better suited for fusion with proteins containing intermolecular disulfide bonds (10). Velamins have the potential to exhibit similar properties to other CaPhs, including safe handling, low cellular toxicity, and minimal background noise. However, what sets velamins apart is their unique ability to naturally emit chemiluminescence at longer wavelengths (λ_max_= 500-508 nm) with no need of a fluorophore to act as a secondary emitter to shift the wavelength. This characteristic makes velamins particularly advantageous for developing novel biotechnological applications, as the green light they produce is less prone to scattering in complex samples compared to the blue light emitted by other CaPhs.

## Materials and Methods

### Collection and molecular identification of *Velamen parallelum* specimens

Specimens of the bioluminescent ctenophore *Velamen parallelum* were collected in May 2017, near Alcatrazes Archipelago (Southeastern Brazil), aboard the Research Vessel MV Alucia (Ocean X). A Triton 3300/3 manned submersible was used to collect the gelatinous animals, which was seen swimming at a depth of around 500-600 m (**Movie 1**). Two specimens were collected using a suction sampler from the submersible and stored in a rotary carousel container during the dives. These specimens were collected under Permit # SISBIO 57721, Brazilian Ministry of the Environment. The samples were immediately preserved and kept in an −80°C freezer until analysis. An aliquot of this material was used for DNA isolation with the DNeasy Mini kit (Qiagen), followed by the amplification of the DNA barcodes 18S rRNA gene (primers CCGAATTCGTCGACAACCTGGTTGATCCTGCCAGT and CCCGGGATCCAAGCTTGATCCTTCTGCAGGTTCACCTAC) (51), and Cytochrome Oxidase subunit 1 - COI (primers GCWGATATGTGTTTACCYMG and TWCCAGAYARRCCWCCAAAAGT) (30). PCR were carried out using the Phusion Plus DNA Polymerase (Invitrogen). DNA amplicons were purified using the QIAquick PCR Purification kit (Qiagen) and sequenced by the Sanger method with an ABI 3730 DNA Analyzer (Applied Biosystems) at the Human Genome and Stem Cell Research Center (CEGH, Brazil). DNA sequences were trimmed by using the software Geneious Prime v. 2023.0.4 (Biomatters Ltd.). 18S and COI sequences from *V. parallelum* of this study were deposited at GenBank database under the accession numbers OQ463859 and PP490733, respectively. 18S and COI sequences from Lobate ctenophores were retrieved from the GenBank database for phylogeny estimation (30) (**Table S4**). Multiple alignments were performed with MUSCLE within Geneious Prime. Bayesian and maximum likelihood phylogenies were estimated from the evolutionary models TN+G+I (18S) and GTR+G+I (COI) with 1000 bootstrap replicates using the IQ-Tree web server (http://iqtree.cibiv.univie.ac.at/). Phylogenies were visualized with FigTree (v1.4.4, tree.bio.ed.ac.uk/software/figtree/). Silhouette images of representative ctenophores were obtained from PhyloPic (www.phylopic.org). Bayesian support values were reported on the phylogeny. *Pleurobrachia bachei* sequences were used as the outgroup.

### Isolation of photoprotein-like sequences from *V. parallelum*

Initially, total RNA was extracted from 100 mg of frozen *V. parallelum* samples using the RNeasy Mini kit (Qiagen), according to the manufacturer’s instructions. Traces of DNA contamination were removed using the DNase I enzyme (Invitrogen). Single-strand cDNA was synthesized with the SuperScriptIV (Invitrogen) from 1 µg of total RNA using oligo(dT)_20_ primers. Two sets of primers were designed from the consensus sequences of mnemiopsin (ATGCCTGGACTGGACGAG and TTAGTATTTGAAGGCGTAGACACCA), and berovin (ATGACTGAACGTCTGAACGAG and TTAGTACTTATAAGCGTAGACTCCG). PCR reactions were carried out using the *V. parallelum* cDNA, primers for mnemiopsin or berovin, and the Q5 High-Fidelity DNA Polymerase (New England Biolabs). DNA amplicons of about 600 bp were purified using the QIAquick PCR Purification kit (Qiagen) and cloned in pGEM-T Easy vector (Promega). A total of 25 positive clones were sequenced by the Sanger method using the universal primers pUC/M13F (CGCCAGGGTTTTCCCAGTCACGAC) and pUC/M13R (CAGGAAACAGCTATGAC), with an ABI 3730 DNA Analyzer (Applied Biosystems) at the Human Genome and Stem Cell Research Center (CEGH, Brazil).

### Expression and purification of recombinant apovelamin

One consensus coding sequence for each cluster of photoproteins (named α-, β- and γ-velamin) was optimized for the recombinant protein expression in *Escherichia coli* and cloned into pET-28a vectors (Novagen) between NdeI and EcoRI restriction sites. N-terminal His-tagged apovelamins were produced by the induction of *E. coli* BL21 (DE3) strain cells with 1 mM IPTG at 15ºC overnight. Cells were harvested by centrifugation and resuspended in a sodium phosphate lysis buffer (50 mM, pH 7.4) containing lysozyme. Recombinant proteins were initially purified by a gravity-flow affinity chromatography using a nickel-charged resin Ni-NTA Agarose (Qiagen) equilibrated with 10 mM imidazole in lysis buffer. A linear gradient of imidazole (100-500 mM) was used to elute the recombinant apovelamins. Protein eluates were analyzed by sodium dodecyl sulfate polyacrylamide gel electrophoresis (SDS-PAGE, 15%). Fractions containing the recombinant apovelamins were pooled and desalted by dialysis in a sodium phosphate buffer (50 mM, pH 7.4). Initial attempts to purify apovelamins by Ni-NTA affinity chromatography frequently resulted in coelution of non-specific proteins.

In order to obtain the purest fractions of each apovelamin, protein-enriched samples were applied onto a gel filtration column (HiLoad 26/600 Superdex 200 pg, Cytiva) connected to a fast protein liquid chromatography (FPLC) system (AKTA pure, Cytiva), previously equilibrated with 25 mM Tris-HCl buffer pH 9. Fractions of 5 mL were collected and assessed for light emission activity. Active fractions were pooled and concentrated using a 3-kDa molecular weight cutoff filter (Vivaspin 500, GE Healthcare), and used for further assays.

Homogeneity was confirmed through SDS-PAGE, indicating the presence of single bands of approximately 25 kDa (Fig. S2). Unless otherwise stated, protein concentrations were determined using a fluorometer Qubit 3.0 (Invitrogen). The theoretical extinction coefficient at 280 nm for each His-tagged apovelamin are: 43,890 M^-1^cm^-1^ (α-velamin), 42,400 M^-1^cm^-1^ (β-velamin) and 43,890 M^-1^cm^-1^ (γ-velamin).

### Light emission assays

Velamin samples were regenerated in 50 mM Tris buffer pH 9 by adding 0.5 µM purified apovelamin, 1 µM coelenterazine (NanoLight) and 10 mM EDTA into 1.5 mL amber microtubes and kept at 4ºC for 12h, unless otherwise stated.

Different times of incubation (0, 2, 4, 8, 10, 12, 16, 22 and 24 hours) were also evaluated for velamin regeneration. Light emission assays were performed using the luminometer Sirius L (Berthold) by injecting 500 µL of 50 mM Tris buffer pH 9 supplemented with 0.5 mM CaCl_2_ to 1 µL of regenerated velamin (Fig. S3). The first 10 seconds of the chemiluminescence assays were considered to estimate the integral of light emission, average and standard deviation from three technical replicates (**Fig. S4**). Activity assays were also performed in different buffers, including sodium phosphate (pH range 6-7.5), Tris-HCl (pH range 7.5-9), and glycine (pH range 9-10). The response of velamins to calcium was evaluated in Tris buffer supplemented with CaCl_2_ concentrations from 10^−2^ to 10^−7^ M. To investigate the thermostability of apovelamins, protein samples were incubated at 25, 37, 50, and 75°C for different periods of time (0, 10, 20, 30, and 60 minutes) in 50 mM Tris buffer pH 9 containing 10 mM EDTA. Then, samples were placed on ice for 30 minutes and regenerated as mentioned above. Additionally, the thermal inactivation of the α-velamin was assessed by incubating the regenerated photoprotein at 37°C for 1, 5, 10, and 15 minutes, followed by the chemiluminescent assay. Photoinactivation of α-velamin was tested by exposing it to visible or UV (395 nm) light for 5, 10, 25 minutes and 1 hour followed by light emission measurements. Light emission spectra for velamins were obtained from the chemiluminescence assays carried out at the Spectrum FLSP 920 photometer composed of two Hamamatsu UV visible R1110 detectors (Edinburgh Instruments, Edinburgh, UK).

### Phylogeny and structural modeling of CaPhs

A multiple sequence alignment was performed using the ten sequences for *V. parallelum* CaPhs, plus hydromedusan and other ctenophore photoproteins, totalizing 45 full-length sequences (**Table S3**) using the Clustal tool (52), with standard parameters. Phylogeny analysis was carried out creating a Maximum Likelihood Tree using MEGA 11 (53) with parameters: Bootstrap method 100, LG with frequencies (+F) model, and Gamma distributed (G) rates among sites. Structure prediction of proteins VparPP2 (velamin-α), VparPP8 (velamin-β), and VparPP5 (velamin-γ) were performed using the software UCSF ChimeraX (v1.6.1, 2023-05-09), which includes an integrated link to the ColabFold-AlphaFold2 suite (54–56). Predictions were carried out for native sequences as well as for His(6)-tagged versions, showing no significant differences. The root-mean-square deviation (RMSD) of atomic positions among all modeled velamin structures using the AlphaFold2 suite (five for each alpha, beta and gamma class, fifteen structures in total) was below 1.3 Å. AlphaFold2 top-ranked predicted structures of VparPPs and other photoprotein models (indicated by the PDB accession number, **Table S3**) were visualized, compared, analyzed and rendered for publication using either UCSF ChimeraX or UCSF Chimera (v1.16) (57).

## Acknowledgments

We thank the Dalio Family Foundation, OceanX, the crew of the M/Y Alucia, and the submarine pilots of the Triton 3300/3 for ctenophore collections in Brazil. We are deeply grateful to Prof. Dr. Marcos Santos of the Oceanographic Institute at the University of São Paulo for organizing and overseeing the Alucia-Brazil expedition. Prof. Dr. Vadim Viviani and Mr. Gabriel Felder Pelentir from the Federal University of São Carlos - Sorocaba, SP, Brazil, for emission spectra experiments of the photoproteins. São Paulo Research Foundation (FAPESP): grants 2017/22501-2 (CVS, AGO); 2019/12605-0 (DMMS); 2022/14964-0 (DMMS); 2020/11189-0 (GGS); 2019/25086-1 (GVM); and the Brazilian National Council for Scientific and Technological (CNPq): 303525/2021-5 (CVS). This study was financed in part by the Coordenação de Aperfeiçoamento de Pessoal de Nível Superior - Brasil (CAPES) - Finance Code 88887.605088/2021-00 (GAG). Yeshiva University Start-up Fund #2022 (AGO).

