## Supporting Information for "Velamins: the first green-light emitting class of wild-type Ca^2+^-regulated photoproteins isolated from the ctenophore *Velamen parallelum*"

[1]Departamento de Química Fundamental, Instituto de Química, Universidade de São Paulo, São Paulo 05508-000, Brazil.
[2]Oceanographic Institute, University of São Paulo, São Paulo, Brazil.
[3]Departamento de Ciências BioMoleculares, Faculdade de Ciências Farmacêuticas de Ribeirão Preto, Universidade de São Paulo, Ribeirão Preto, Brazil.
[4]Department of Chemistry and Biochemistry, Yeshiva University, New York, USA.
[5]Center for Marine Biology, University of São Paulo, São Sebastião, Brazil.
[6]Department of Natural Sciences, Baruch College, City University of New York, New York, NY, 10010, USA.
[7]Department of Ichthyology, Division of Vertebrate Zoology, American Museum of Natural History, New York, NY 10024, USA.
[8]The John B. Pierce Laboratory, New Haven, CT 06519, USA.
[9]Department of Cellular and Molecular Physiology, Yale School of Medicine, New Haven, Connecticut, USA.
*Anderson G. Oliveira.

**Supporting Information**

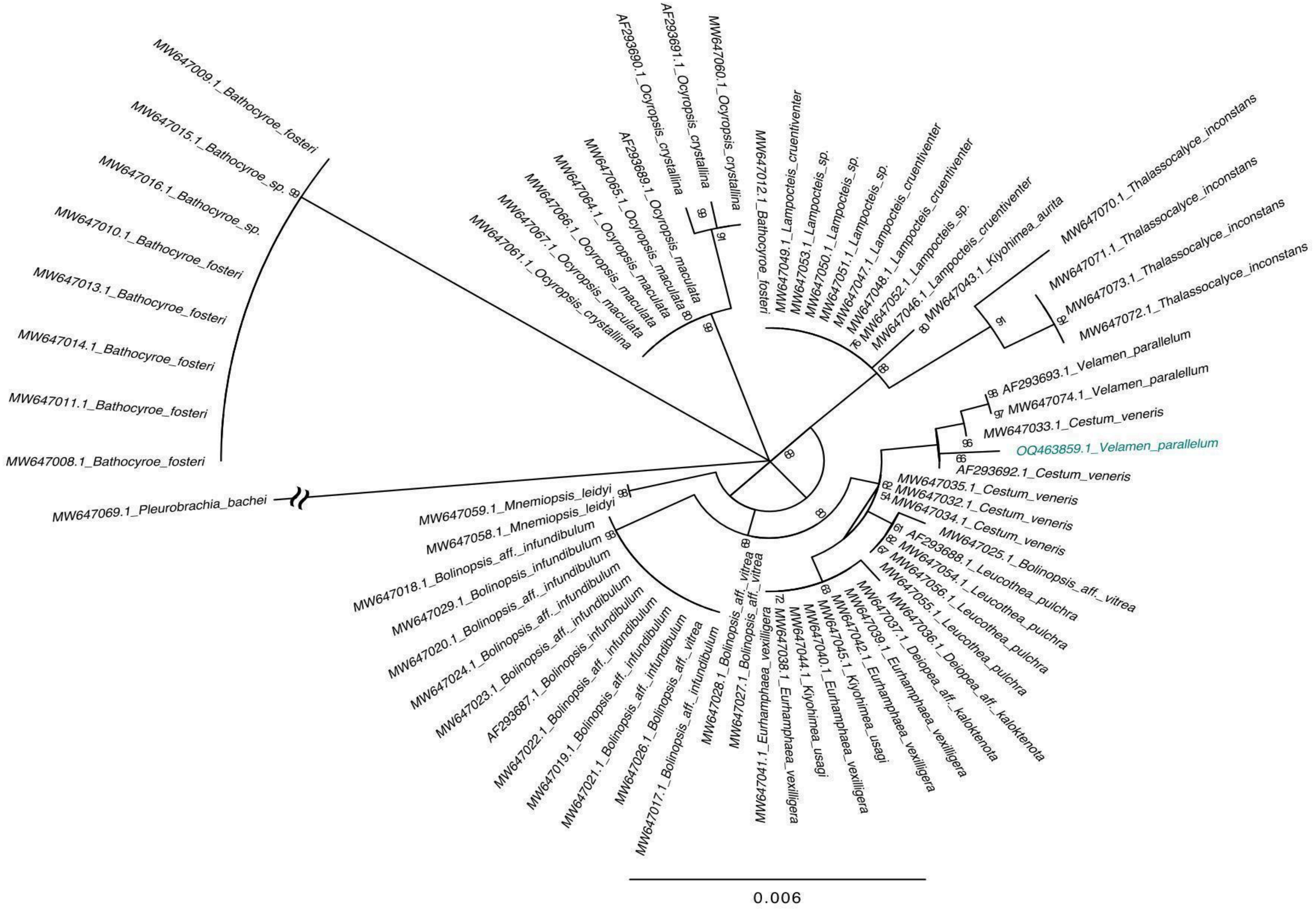


**Fig. S1.** Maximum likelihood estimated phylogeny of Lobate ctenophores based on 18S rDNA gene fragments. Bootstrap support values >50% are shown. DNA sequence from the *Velamen parallelum* specimen under analysis in this study is highlighted.

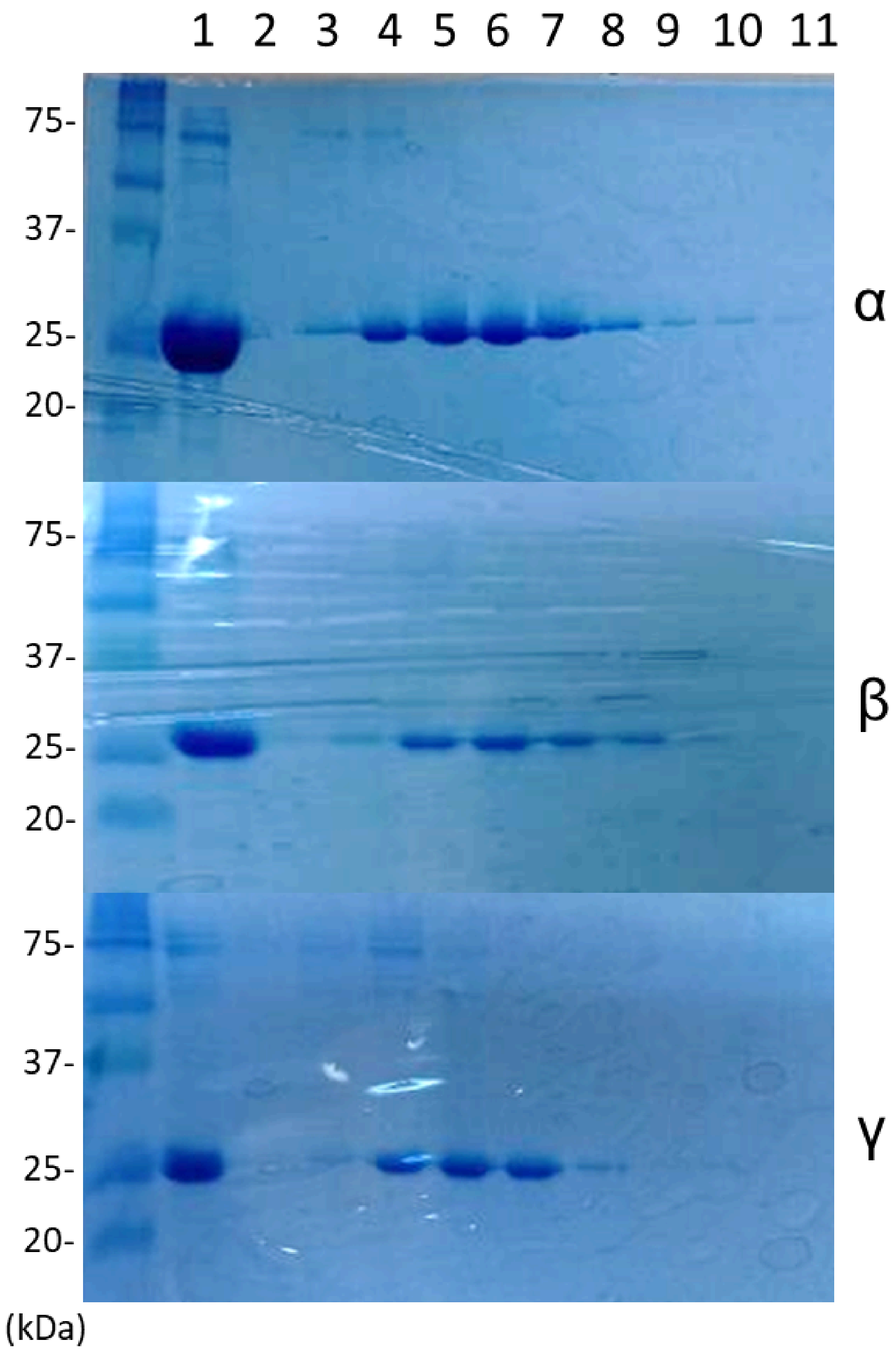


**Fig. S2.** SDS-PAGE gel electrophoresis of α-, β- and γ-velamins purified by gravity-flow chromatography. Total protein fraction (1), followed by elution fractions with 0.1 M (2 and 3); 0.2 M (4 and 5); 0.3 M (6 and 7); 0.4 M (8 and 9); and 0.5 M (10 and 11) imidazole.

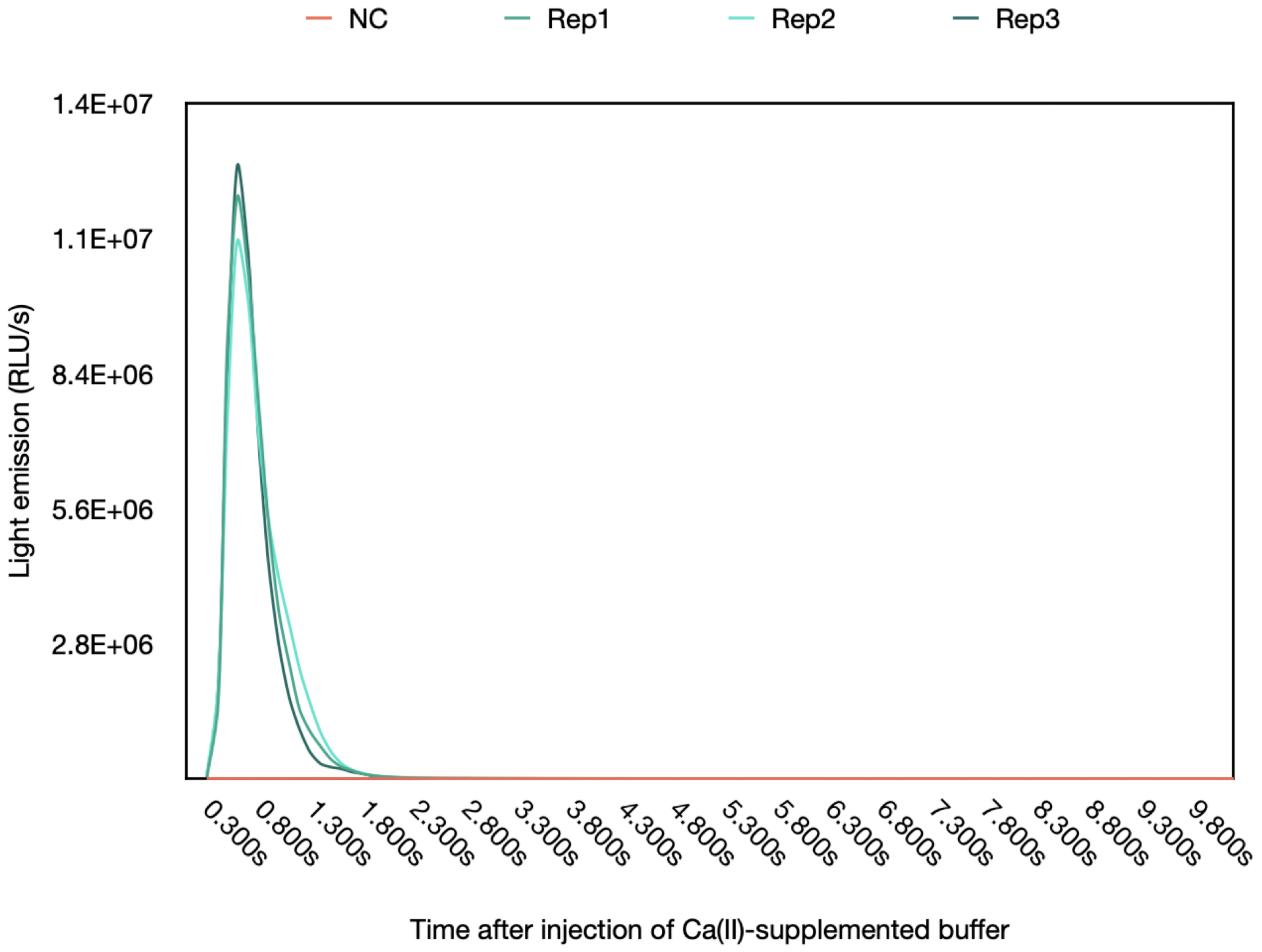


**Fig. S3.** Example of raw data obtained from a luminometer (Berthold) in a light emission assay using α-velamin. 500 µL of 50 mM Tris buffer pH 9 supplemented with 0.5 mM $CaCl_2$ were injected to test tubes containing 1 µL of α-velamin regenerated at 4ºC during 8h in 1.5 mL amber microtubes using 50 mM Tris buffer pH 9, 0.5 µM purified apo-α-velamin, 1 µM coelenterazine (NanoLight) and 10 mM EDTA. Three technical replicates and a negative control were assessed.

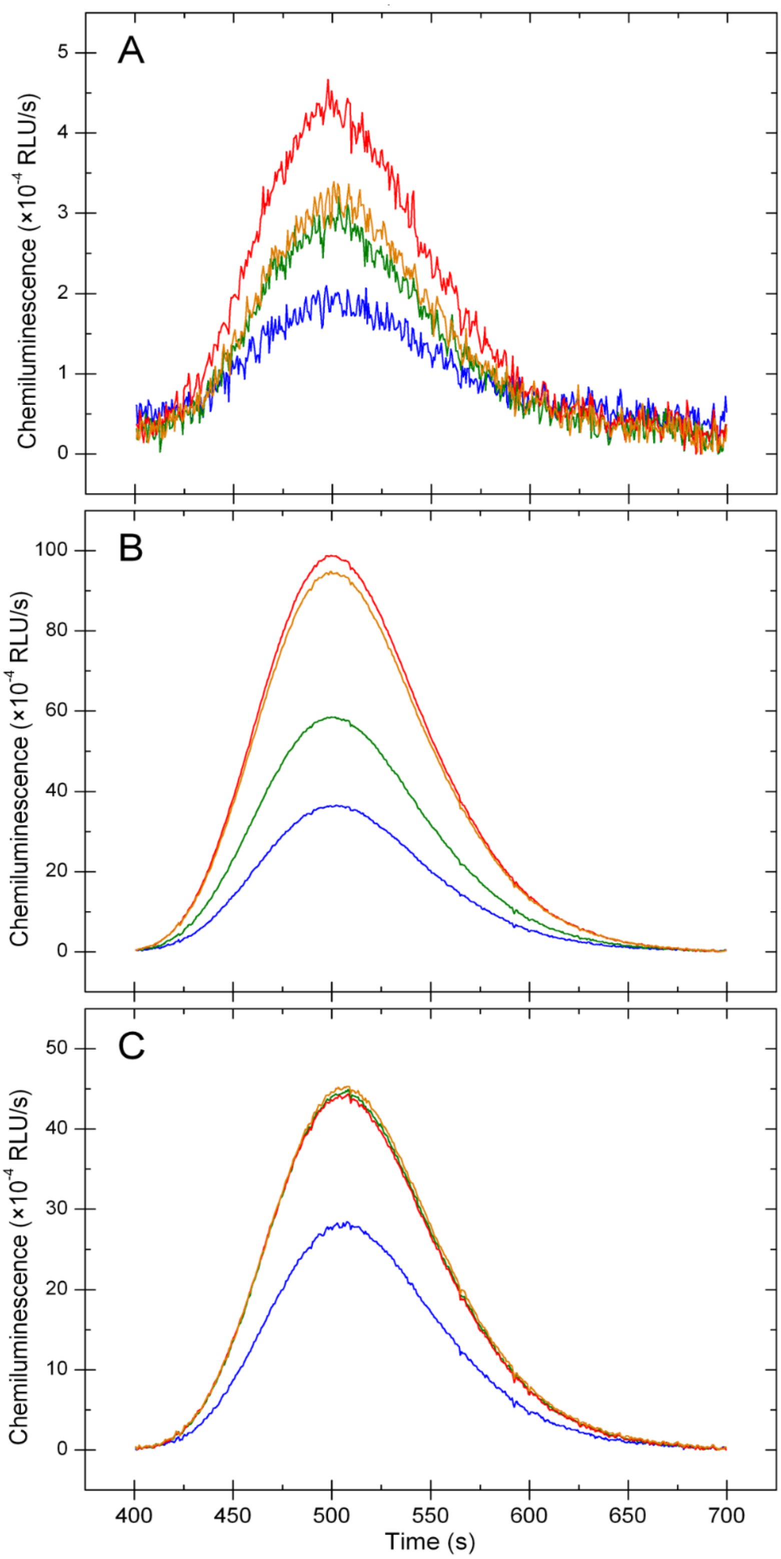


**Fig. S4.** Light emission spectra obtained from α- (A), β- (B) and γ-velamins (C) in different pH (–7, –7.5, –8 and –9). Assays were carried out at the Spectrum FLSP 920 photometer composed of two Hamamatsu UV visible R1110 detectors (Edinburgh Instruments) in cuvettes containing 50 μL of velamins regenerated at 4ºC during 16h in 1.5 mL amber microtubes using 50 mM Tris buffer pH 9, 0.5 μM purified apovelamin, 1 μM coelenterazine (NanoLight) and 10 mM EDTA. Light emission was recorded after adding 1 mL of 50 mM Tris buffer pH 9 supplemented with 0.5 mM $CaCl_2$.

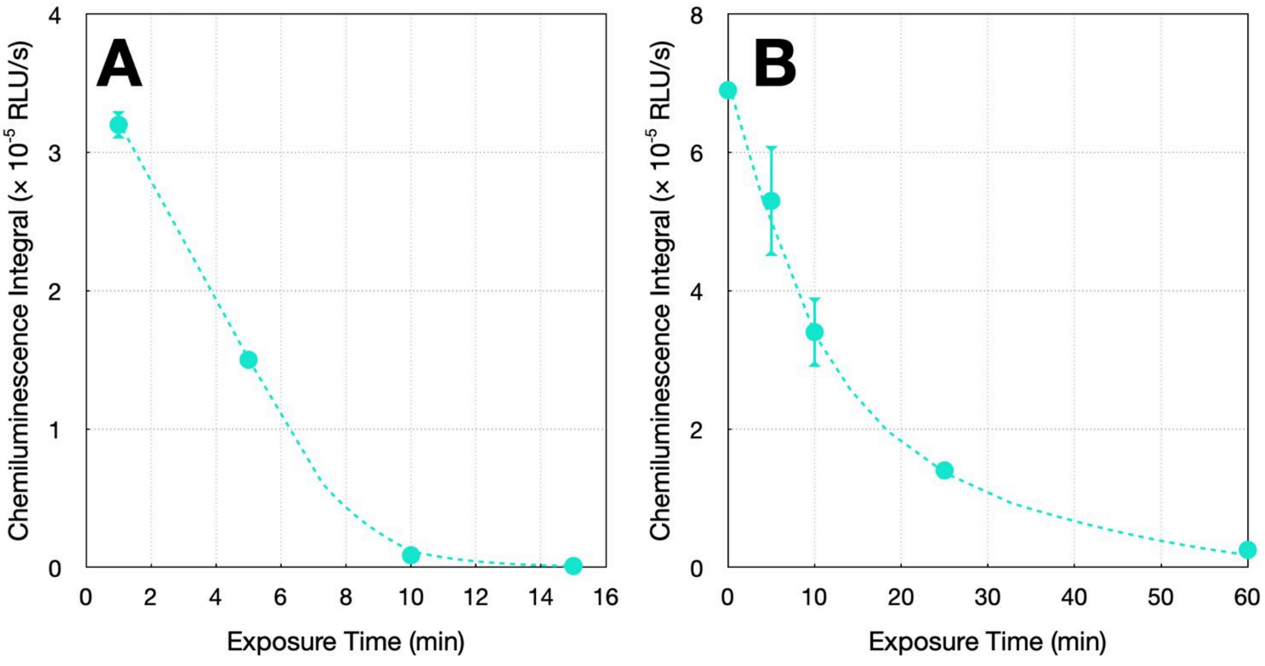


**Fig. S5.** (**A**) Thermal inactivation of α-velamin incubated at 37ºC for 1, 5, 10, and 15 minutes. (**B**) Photoinactivation assay using α-velamin exposed to visible light for 5, 10, 25 minutes and 1 hour. In both experiments, α-velamin was regenerated by using 50 mM Tris buffer pH 9, 0.5 μM purified apo-α-velamin, 1 μM coelenterazine (NanoLight) and 10 mM EDTA at 4ºC during 12h in 1.5 mL amber microtubes. After thermal or photoinactivation experiments, 500 μL of 50 mM Tris buffer pH 9 supplemented with 0.5 mM $CaCl_2$ were injected to test tubes containing 1 μL of α-velamin and the light emission was recorded in a luminometer (Berthold). Three technical replicates were assessed. Average and standard deviation are calculated from the estimation of integral values obtained for technical replicates.

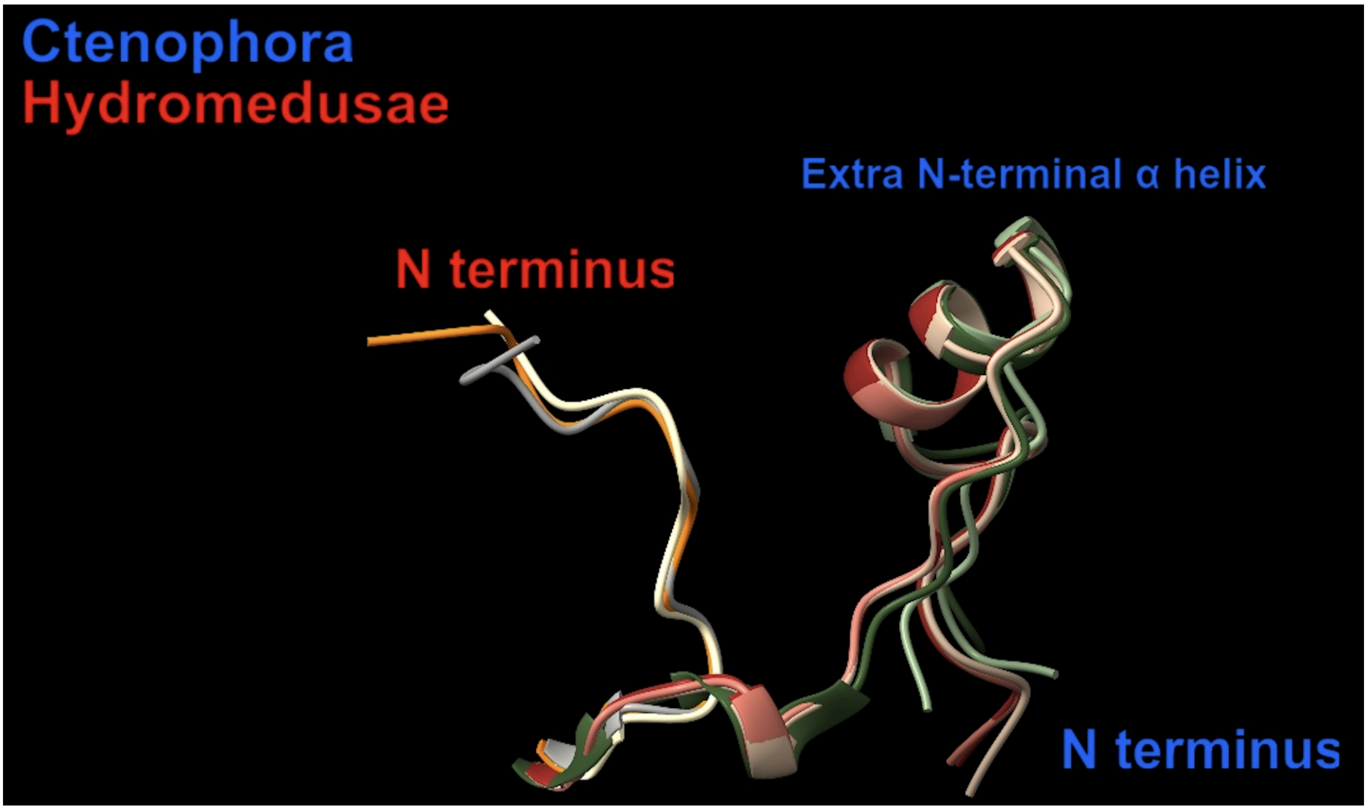


**Fig. S6.** Structural comparison at the N-termini of CaPhs from ctenophores and hydromedusae. More details at **Movie 3**. Structure prediction of proteins VparPP2 (velamin-α), VparPP8 (velamin-β), and VparPP5 (velamin-γ) were performed using the software UCSF ChimeraX, which includes an integrated link to the ColabFold-AlphaFold2 suite (1–3). AlphaFold2 top-ranked predicted structures of VparPPs and other photoprotein models (indicated by name and PDB accession number, **Table S3**) were visualized, compared, analyzed and rendered for publication using either UCSF ChimeraX.

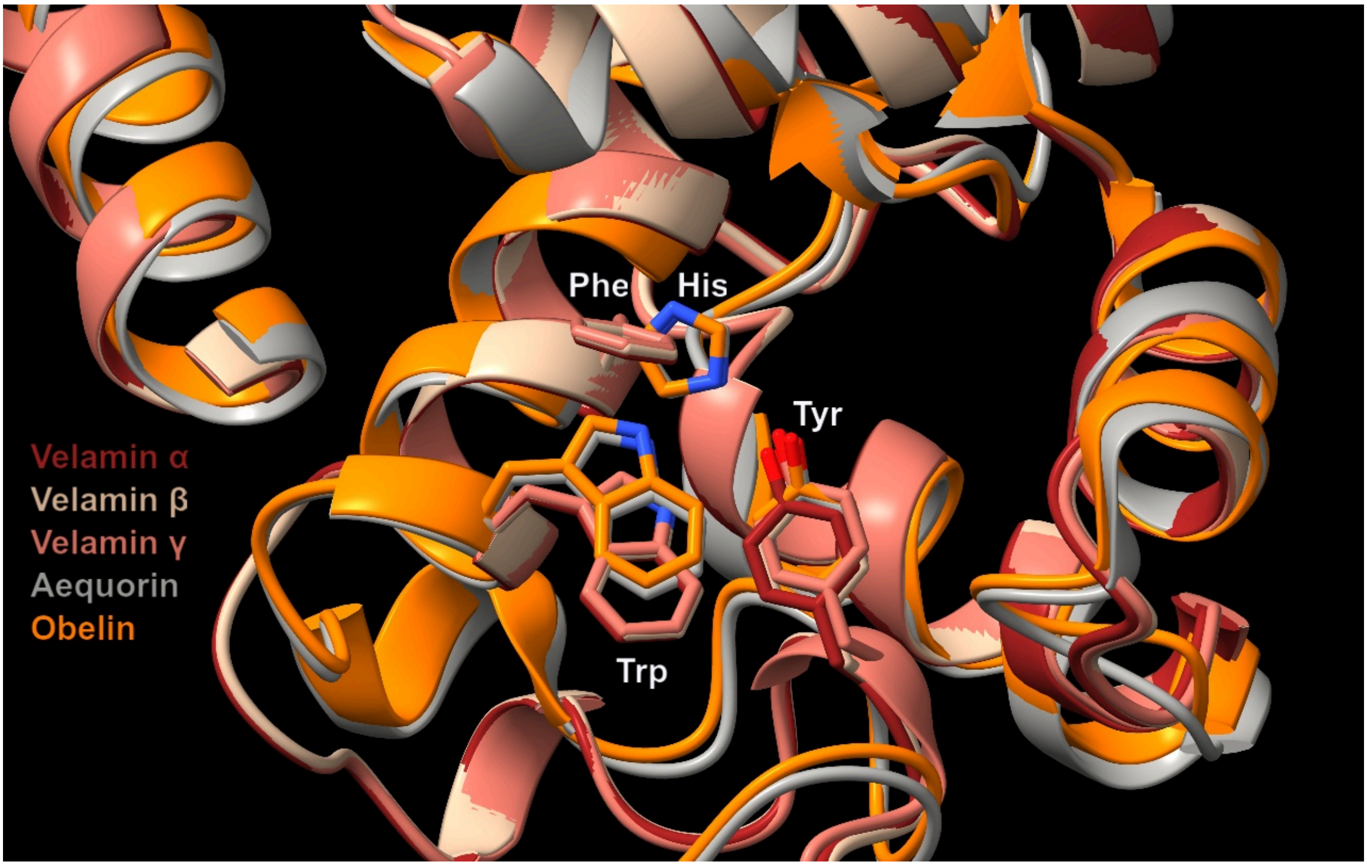


**Fig. S7**. Superimposed predicted structures from photoproteins highlighting the replacement of the conserved triad His-Trp-Tyr, found in hydromedusan CaPhs, with the equivalent Phe-Trp-Tyr in ctenophore CaPhs. More details at **Movie 4**. Structure prediction of proteins VparPP2 (velamin-α), VparPP8 (velamin-β), and VparPP5 (velamin-γ) were performed using the software UCSF ChimeraX, which includes an integrated link to the ColabFold-AlphaFold2 suite (1–3). AlphaFold2 top-ranked predicted structures of VparPPs and other photoprotein models (indicated by name and PDB accession number, **Table S3**) were visualized, compared, analyzed and rendered for publication using either UCSF ChimeraX.

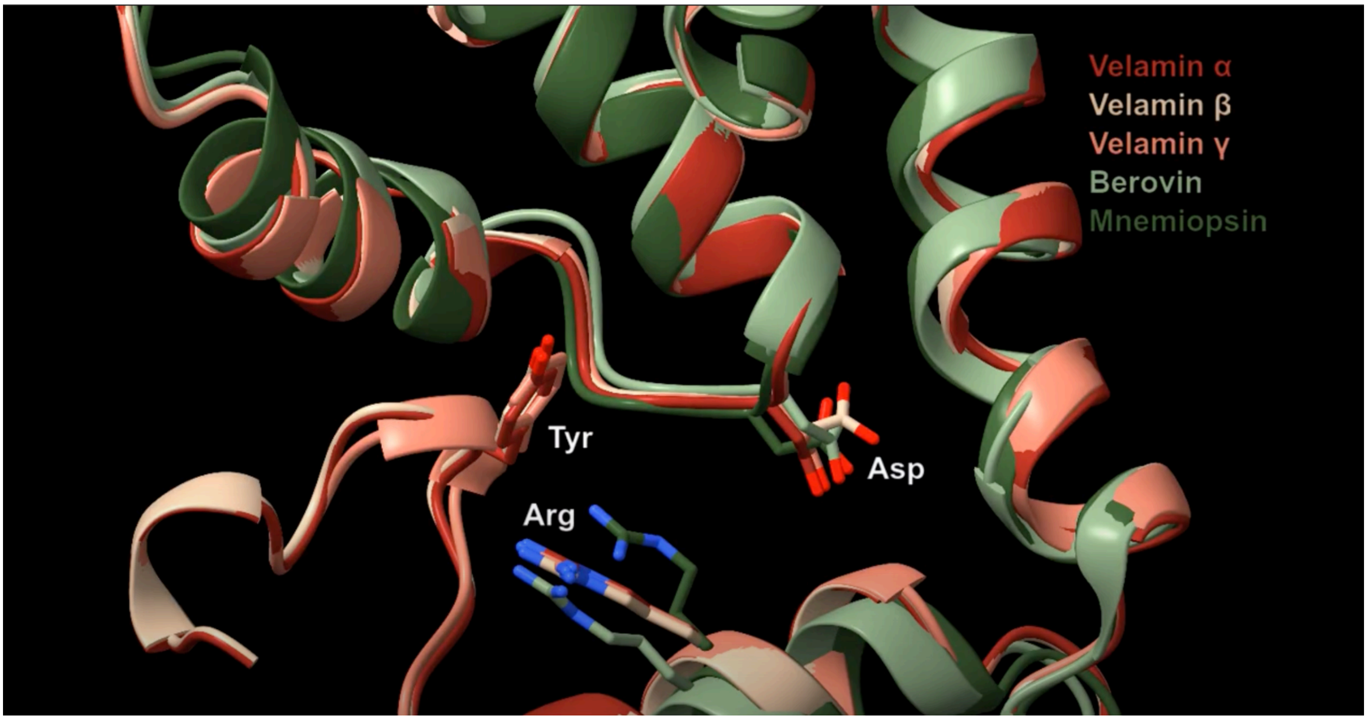


**Fig. S8.** Conserved triad Arg40-Asp157-Tyr203 (VparPP2) proposed to be involved in light emission in ctenophore CaPhs. More details at **Movie 5**. Structure prediction of proteins VparPP2 (velamin-α), VparPP8 (velamin-β), and VparPP5 (velamin-γ) were performed using the software UCSF ChimeraX, which includes an integrated link to the ColabFold-AlphaFold2 suite (1–3). AlphaFold2 top-ranked predicted structures of VparPPs and other photoprotein models (indicated by name and PDB accession number, **Table S3**) were visualized, compared, analyzed and rendered for publication using either UCSF ChimeraX.

**Table S1.** Percentage of identity among CaPhs from hydromedusae and ctenophores[a]. *V. parallelum* CaPhs are highlighted.

| | AEQ1_AEQVI | AEQ2_AEQVI | AEQ_CO | AEQ_MA | AEQ_PA | Berovin | Bolinopsin | Bolinopsin_gr | **VparPP1** | **VparPP2** | **VparPP3** | **VparPP4** | **VparPP5** | **VparPP6** | **VparPP7** | **VparPP8** | **VparPP9** | **VparPP10** |
|---|---|---|---|---|---|---|---|---|---|---|---|---|---|---|---|---|---|---|
| AEQ1_AEQVI | | 90.3 | 83.6 | 85.1 | 83.6 | 24.9 | 24.4 | 23.9 | 23.4 | 23.4 | 23.9 | 23.9 | 24.9 | 24.4 | 24.4 | 24.9 | 23.4 | 23.4 |
| AEQ2_AEQVI | 90.3 | | 80.0 | 82.1 | 83.6 | 25.9 | 23.9 | 24.4 | 23.4 | 23.4 | 23.9 | 23.4 | 23.9 | 23.9 | 23.9 | 24.4 | 23.4 | 23.4 |
| AEQ_CO | 83.6 | 80.0 | | 82.6 | 83.1 | 24.0 | 24.0 | 24.0 | 24.0 | 24.0 | 24.5 | 23.0 | 24.5 | 23.5 | 24.5 | 24.5 | 23.5 | 24.0 |
| AEQ_MA | 85.1 | 82.1 | 82.6 | | 93.8 | 25.0 | 24.5 | 25.0 | 24.0 | 24.0 | 24.5 | 23.5 | 23.5 | 24.0 | 24.5 | 24.5 | 24.0 | 24.0 |
| AEQ_PA | 83.6 | 83.6 | 83.1 | 93.8 | | 23.0 | 23.0 | 23.0 | 23.0 | 23.0 | 23.5 | 21.9 | 22.4 | 22.4 | 23.0 | 23.0 | 22.4 | 23.0 |
| Berovin | 24.9 | 25.9 | 24.0 | 25.0 | 23.0 | | 84.6 | 88.5 | 87.0 | 86.1 | 87.5 | 89.9 | 86.5 | 90.4 | 86.5 | 90.4 | 88.0 | 87.0 |
| Bolinopsin | 24.4 | 23.9 | 24.0 | 24.5 | 23.0 | 84.6 | | 85.0 | 84.5 | 84.5 | 85.0 | 86.5 | 84.1 | 87.0 | 84.1 | 86.5 | 84.5 | 84.5 |
| Bolinopsin_gr | 23.9 | 24.4 | 24.0 | 25.0 | 23.0 | 88.5 | 85.0 | | 85.0 | 84.1 | 85.5 | 88.9 | 91.8 | 89.4 | 90.8 | 88.9 | 85.5 | 85.0 |
| **VparPP1** | 23.4 | 23.4 | 24.0 | 24.0 | 23.0 | 87.0 | 84.5 | 85.0 | | 98.6 | 99.5 | 91.8 | 87.0 | 91.3 | 90.8 | 91.3 | 97.1 | 99.5 |
| **VparPP2** | 23.4 | 23.4 | 24.0 | 24.0 | 23.0 | 86.1 | 84.5 | 84.1 | 98.6 | | 98.1 | 90.8 | 86.0 | 90.3 | 89.9 | 90.3 | 96.1 | 99.0 |
| **VparPP3** | 23.9 | 23.9 | 24.5 | 24.5 | 23.5 | 87.5 | 85.0 | 85.5 | 99.5 | 98.1 | | 92.3 | 87.4 | 91.8 | 91.3 | 91.8 | 96.6 | 99.0 |
| **VparPP4** | 23.9 | 23.4 | 23.0 | 23.5 | 21.9 | 89.9 | 86.5 | 88.9 | 91.8 | 90.8 | 92.3 | | 88.9 | 99.5 | 89.4 | 98.6 | 94.7 | 91.8 |
| **VparPP5** | 24.9 | 23.9 | 24.5 | 23.5 | 22.4 | 86.5 | 84.1 | 91.8 | 87.0 | 86.0 | 87.4 | 88.9 | | 89.4 | 96.1 | 89.4 | 86.0 | 87.0 |
| **VparPP6** | 24.4 | 23.9 | 23.5 | 24.0 | 22.4 | 90.4 | 87.0 | 89.4 | 91.3 | 90.3 | 91.8 | 99.5 | 89.4 | | 89.9 | 99.0 | 94.2 | 91.3 |
| **VparPP7** | 24.4 | 23.9 | 24.5 | 24.5 | 23.0 | 86.5 | 84.1 | 90.8 | 90.8 | 89.9 | 91.3 | 89.4 | 96.1 | 89.9 | | 89.9 | 89.9 | 90.8 |
| **VparPP8** | 24.9 | 24.4 | 24.5 | 24.5 | 23.0 | 90.4 | 86.5 | 88.9 | 91.3 | 90.3 | 91.8 | 98.6 | 89.4 | 99.0 | 89.9 | | 93.2 | 91.3 |
| **VparPP9** | 23.4 | 23.4 | 23.5 | 24.0 | 22.4 | 88.0 | 84.5 | 85.5 | 97.1 | 96.1 | 96.6 | 94.7 | 86.0 | 94.2 | 89.9 | 93.2 | | 97.1 |
| **VparPP10** | 23.4 | 23.4 | 24.0 | 24.0 | 23.0 | 87.0 | 84.5 | 85.0 | 99.5 | 99.0 | 99.0 | 91.8 | 87.0 | 91.3 | 90.8 | 91.3 | 97.1 | |
| Clytin1 | 60.7 | 58.7 | 61.0 | 60.5 | 57.9 | 25.1 | 26.1 | 25.6 | 25.1 | 25.1 | 25.6 | 24.6 | 25.1 | 25.1 | 26.1 | 25.1 | 24.6 | 25.1 |
| Clytin2 | 61.7 | 58.7 | 61.0 | 60.0 | 60.0 | 24.1 | 24.1 | 23.7 | 22.8 | 22.8 | 23.3 | 22.8 | 23.7 | 23.3 | 24.1 | 23.3 | 22.4 | 22.8 |
| Clytin3 | 63.3 | 61.2 | 63.6 | 61.0 | 60.0 | 25.1 | 26.6 | 25.1 | 25.1 | 25.1 | 25.6 | 26.1 | 26.6 | 26.6 | 26.1 | 27.1 | 25.6 | 25.1 |
| Clytin-I | 62.8 | 60.7 | 60.5 | 62.1 | 60.5 | 25.6 | 26.6 | 26.1 | 25.6 | 25.6 | 26.1 | 25.1 | 25.6 | 25.6 | 26.6 | 25.6 | 25.1 | 25.6 |
| MC1 | 68.7 | 65.1 | 72.8 | 67.7 | 67.2 | 23.5 | 24.0 | 22.4 | 22.4 | 22.4 | 23.0 | 23.5 | 23.5 | 23.5 | 23.0 | 24.0 | 23.0 | 22.4 |
| MC2 | 68.2 | 65.1 | 71.8 | 67.2 | 66.7 | 23.5 | 24.0 | 22.4 | 22.4 | 22.4 | 23.0 | 23.5 | 23.5 | 23.5 | 23.0 | 24.0 | 23.0 | 22.4 |
| MC3 | 68.7 | 65.1 | 72.3 | 67.7 | 67.2 | 23.5 | 24.0 | 22.4 | 22.4 | 22.4 | 23.0 | 23.5 | 23.5 | 23.5 | 23.0 | 24.0 | 23.0 | 22.4 |
| MC4 | 68.2 | 64.6 | 71.8 | 67.2 | 66.7 | 23.0 | 24.0 | 21.9 | 22.4 | 22.4 | 22.4 | 23.0 | 23.5 | 23.0 | 22.4 | 23.5 | 23.0 | 22.4 |
| MC5 | 67.7 | 64.1 | 71.3 | 66.7 | 66.7 | 23.0 | 23.5 | 21.9 | 21.9 | 21.9 | 22.4 | 23.0 | 23.0 | 23.0 | 22.4 | 23.5 | 22.4 | 21.9 |
| MI17 | 70.3 | 66.7 | 72.8 | 68.7 | 68.2 | 23.5 | 24.0 | 22.4 | 22.4 | 22.4 | 23.0 | 23.5 | 23.5 | 23.5 | 23.0 | 24.0 | 23.0 | 22.4 |
| MleiPP1 | 23.4 | 24.4 | 24.0 | 24.0 | 22.4 | 88.5 | 86.4 | 87.0 | 93.2 | 93.7 | 92.8 | 90.3 | 86.5 | 89.9 | 88.9 | 89.9 | 92.8 | 93.7 |
| MleiPP2 | 23.4 | 24.4 | 24.0 | 24.0 | 23.0 | 91.3 | 86.0 | 90.3 | 89.9 | 89.4 | 90.3 | 93.7 | 90.8 | 94.2 | 90.8 | 94.2 | 89.4 | 90.3 |
| MleiPP3 | 22.8 | 23.9 | 23.5 | 23.5 | 22.4 | 91.3 | 86.5 | 89.4 | 91.8 | 91.3 | 91.3 | 93.7 | 88.9 | 93.2 | 88.9 | 93.2 | 91.3 | 92.3 |

| | | | | | | | | | | | | | | | | | | |
|---|---|---|---|---|---|---|---|---|---|---|---|---|---|---|---|---|---|---|
| MleiPP4 | 23.4 | 23.9 | 24.0 | 24.0 | 23.0 | 88.0 | 85.4 | 86.0 | 92.8 | 93.2 | 92.3 | 89.9 | 86.0 | 89.4 | 88.4 | 89.4 | 92.3 | 93.2 |
| MleiPP5 | 23.4 | 24.4 | 24.0 | 24.0 | 23.0 | 90.9 | 85.5 | 89.4 | 89.9 | 89.4 | 90.3 | 93.2 | 90.3 | 93.7 | 90.3 | 93.7 | 89.4 | 90.3 |
| MleiPP6 | 23.4 | 24.4 | 23.5 | 24.0 | 22.4 | 88.0 | 87.4 | 87.0 | 93.2 | 93.7 | 92.8 | 89.9 | 87.4 | 89.4 | 89.9 | 89.4 | 92.3 | 93.7 |
| MleiPP7 | 22.3 | 23.4 | 23.0 | 23.0 | 21.9 | 90.4 | 85.5 | 88.9 | 91.3 | 90.8 | 90.8 | 92.8 | 88.9 | 92.3 | 88.9 | 92.3 | 90.8 | 91.8 |
| MleiPP8 | 25.4 | 24.9 | 24.5 | 24.0 | 23.0 | 87.5 | 85.0 | 90.3 | 87.0 | 86.0 | 87.4 | 88.4 | 96.1 | 88.9 | 93.7 | 88.9 | 86.0 | 87.0 |
| MleiPP9 | 25.4 | 24.9 | 24.5 | 24.0 | 23.0 | 87.0 | 84.5 | 89.9 | 86.5 | 85.5 | 87.0 | 87.9 | 95.7 | 88.4 | 93.2 | 88.4 | 85.5 | 86.5 |
| MleiPP10 | 25.4 | 24.9 | 24.5 | 24.0 | 23.0 | 87.0 | 84.5 | 89.9 | 86.5 | 85.5 | 87.0 | 87.9 | 95.7 | 88.4 | 93.2 | 88.4 | 85.5 | 86.5 |
| OBL | 6.5 | 6.5 | 7.5 | 8.0 | 7.5 | 9.2 | 9.2 | 9.6 | 10.0 | 10.0 | 10.0 | 9.6 | 8.7 | 9.6 | 9.6 | 9.6 | 10.0 | 10.0 |
| OG1 | 62.6 | 64.1 | 64.6 | 64.1 | 62.6 | 26.0 | 26.0 | 25.0 | 25.5 | 25.5 | 26.0 | 25.5 | 25.0 | 26.0 | 25.5 | 26.0 | 26.0 | 25.5 |
| OG2 | 62.1 | 63.6 | 64.1 | 63.6 | 62.1 | 26.0 | 26.0 | 25.0 | 25.5 | 25.5 | 26.0 | 25.5 | 25.0 | 26.0 | 25.5 | 26.0 | 26.0 | 25.5 |

**Table S1. Cont.** Percentage of identity among CaPhs from hydromedusae and ctenophores[a]. *V. parallelum* CaPhs are highlighted.

| | Clytin1 | Clytin2 | Clytin3 | Clytin-I | MC1 | MC2 | MC3 | MC4 | MC5 | MI17 | MleiPP1 | MleiPP2 | MleiPP3 | MleiPP4 | MleiPP5 | MleiPP6 | MleiPP7 | MleiPP8 | MleiPP9 | MleiPP10 | OBL | OG1 | OG2 |
|---|---|---|---|---|---|---|---|---|---|---|---|---|---|---|---|---|---|---|---|---|---|---|---|
| AEQ1_AEQVI | 60.7 | 61.7 | 63.3 | 62.8 | 68.7 | 68.2 | 68.7 | 68.2 | 67.7 | 70.3 | 23.4 | 23.4 | 22.8 | 23.4 | 23.4 | 23.4 | 22.3 | 25.4 | 25.4 | 25.4 | 6.5 | 62.6 | 62.1 |
| AEQ2_AEQVI | 58.7 | 58.7 | 61.2 | 60.7 | 65.1 | 65.1 | 65.1 | 64.6 | 64.1 | 66.7 | 24.4 | 24.4 | 23.9 | 23.9 | 24.4 | 24.4 | 23.4 | 24.9 | 24.9 | 24.9 | 6.5 | 64.1 | 63.6 |
| AEQ_CO | 61.0 | 61.0 | 63.6 | 60.5 | 72.8 | 71.8 | 72.3 | 71.8 | 71.3 | 72.8 | 24.0 | 24.0 | 23.5 | 24.0 | 24.0 | 23.5 | 23.0 | 24.5 | 24.5 | 24.5 | 7.5 | 64.6 | 64.1 |
| AEQ_MA | 60.5 | 60.0 | 61.0 | 62.1 | 67.7 | 67.2 | 67.7 | 67.2 | 66.7 | 68.7 | 24.0 | 24.0 | 23.5 | 24.0 | 24.0 | 24.0 | 23.0 | 24.0 | 24.0 | 24.0 | 8.0 | 64.1 | 63.6 |
| AEQ_PA | 57.9 | 60.0 | 60.0 | 60.5 | 67.2 | 66.7 | 67.2 | 66.7 | 66.7 | 68.2 | 22.4 | 23.0 | 22.4 | 23.0 | 23.0 | 22.4 | 21.9 | 23.0 | 23.0 | 23.0 | 7.5 | 62.6 | 62.1 |
| Berovin | 25.1 | 24.1 | 25.1 | 25.6 | 23.5 | 23.5 | 23.5 | 23.0 | 23.0 | 23.5 | 88.5 | 91.3 | 91.3 | 88.0 | 90.9 | 88.0 | 90.4 | 87.5 | 87.0 | 87.0 | 9.2 | 26.0 | 26.0 |
| Bolinopsin | 26.1 | 24.1 | 26.6 | 26.6 | 24.0 | 24.0 | 24.0 | 24.0 | 23.5 | 24.0 | 86.4 | 86.0 | 86.5 | 85.4 | 85.5 | 87.4 | 85.5 | 85.0 | 84.5 | 84.5 | 9.2 | 26.0 | 26.0 |
| Bolinopsin_gr | 25.6 | 23.7 | 25.1 | 26.1 | 22.4 | 22.4 | 22.4 | 21.9 | 21.9 | 22.4 | 87.0 | 90.3 | 89.4 | 86.0 | 89.4 | 87.0 | 88.9 | 90.3 | 89.9 | 89.9 | 9.6 | 25.0 | 25.0 |
| **VparPP1** | 25.1 | 22.8 | 25.1 | 25.6 | 22.4 | 22.4 | 22.4 | 22.4 | 21.9 | 22.4 | 93.2 | 89.9 | 91.8 | 92.8 | 89.9 | 93.2 | 91.3 | 87.0 | 86.5 | 86.5 | 10.0 | 25.5 | 25.5 |
| **VparPP2** | 25.1 | 22.8 | 25.1 | 25.6 | 22.4 | 22.4 | 22.4 | 22.4 | 21.9 | 22.4 | 93.7 | 89.4 | 91.3 | 93.2 | 89.4 | 93.7 | 90.8 | 86.0 | 85.5 | 85.5 | 10.0 | 25.5 | 25.5 |
| **VparPP3** | 25.6 | 23.3 | 25.6 | 26.1 | 23.0 | 23.0 | 23.0 | 22.4 | 22.4 | 23.0 | 92.8 | 90.3 | 91.3 | 92.3 | 90.3 | 92.8 | 90.8 | 87.4 | 87.0 | 87.0 | 10.0 | 26.0 | 26.0 |
| **VparPP4** | 24.6 | 22.8 | 26.1 | 25.1 | 23.5 | 23.5 | 23.5 | 23.0 | 23.0 | 23.5 | 90.3 | 93.7 | 93.7 | 89.9 | 93.2 | 89.9 | 92.8 | 88.4 | 87.9 | 87.9 | 9.6 | 25.5 | 25.5 |
| **VparPP5** | 25.1 | 23.7 | 26.6 | 25.6 | 23.5 | 23.5 | 23.5 | 23.5 | 23.0 | 23.5 | 86.5 | 90.8 | 88.9 | 86.0 | 90.3 | 87.4 | 88.9 | 96.1 | 95.7 | 95.7 | 8.7 | 25.0 | 25.0 |
| **VparPP6** | 25.1 | 23.3 | 26.6 | 25.6 | 23.5 | 23.5 | 23.5 | 23.0 | 23.0 | 23.5 | 89.9 | 94.2 | 93.2 | 89.4 | 93.7 | 89.4 | 92.3 | 88.9 | 88.4 | 88.4 | 9.6 | 26.0 | 26.0 |
| **VparPP7** | 26.1 | 24.1 | 26.1 | 26.6 | 23.0 | 23.0 | 23.0 | 22.4 | 22.4 | 23.0 | 88.9 | 90.8 | 88.9 | 88.4 | 90.3 | 89.9 | 88.9 | 93.7 | 93.2 | 93.2 | 9.6 | 25.5 | 25.5 |
| **VparPP8** | 25.1 | 23.3 | 27.1 | 25.6 | 24.0 | 24.0 | 24.0 | 23.5 | 23.5 | 24.0 | 89.9 | 94.2 | 93.2 | 89.4 | 93.7 | 89.4 | 92.3 | 88.9 | 88.4 | 88.4 | 9.6 | 26.0 | 26.0 |
| **VparPP9** | 24.6 | 22.4 | 25.6 | 25.1 | 23.0 | 23.0 | 23.0 | 23.0 | 22.4 | 23.0 | 92.8 | 89.4 | 91.3 | 92.3 | 89.4 | 92.3 | 90.8 | 86.0 | 85.5 | 85.5 | 10.0 | 26.0 | 26.0 |
| **VparPP10** | 25.1 | 22.8 | 25.1 | 25.6 | 22.4 | 22.4 | 22.4 | 22.4 | 21.9 | 22.4 | 93.7 | 90.3 | 92.3 | 93.2 | 90.3 | 93.7 | 91.8 | 87.0 | 86.5 | 86.5 | 10.0 | 25.5 | 25.5 |
| Clytin1 | | 87.4 | 81.3 | 92.9 | 60.0 | 59.5 | 59.5 | 59.0 | 58.5 | 59.5 | 25.6 | 25.6 | 24.6 | 25.1 | 25.6 | 25.1 | 24.1 | 25.6 | 25.6 | 25.6 | 8.3 | 77.9 | 77.4 |
| Clytin2 | 87.4 | | 82.8 | 86.9 | 61.0 | 60.5 | 60.5 | 60.0 | 60.0 | 59.5 | 23.3 | 23.7 | 22.4 | 22.8 | 23.7 | 22.8 | 22.0 | 24.1 | 24.1 | 24.1 | 7.6 | 74.4 | 73.8 |
| Clytin3 | 81.3 | 82.8 | | 82.3 | 64.6 | 64.6 | 64.6 | 64.1 | 63.6 | 63.6 | 25.1 | 25.1 | 24.1 | 24.6 | 25.1 | 24.6 | 23.6 | 26.6 | 26.6 | 26.6 | 6.9 | 75.4 | 74.9 |

| | | | | | | | | | | | | | | | | | | | | | | | |
|---|---|---|---|---|---|---|---|---|---|---|---|---|---|---|---|---|---|---|---|---|---|---|---|
| Clytin-I | 92.9 | 86.9 | 82.3 | | 59.5 | 59.5 | 59.5 | 59.0 | 59.0 | 60.0 | 26.1 | 26.1 | 25.1 | 25.6 | 26.1 | 25.6 | 24.6 | 26.1 | 26.1 | 26.1 | 8.3 | 76.4 | 75.9 |
| MC1 | 60.0 | 61.0 | 64.6 | 59.5 | | 99.0 | 99.5 | 98.5 | 98.5 | 98.0 | 22.4 | 22.4 | 22.4 | 23.0 | 22.4 | 23.0 | 21.9 | 23.5 | 23.5 | 23.5 | 6.0 | 64.6 | 64.1 |
| MC2 | 59.5 | 60.5 | 64.6 | 59.5 | 99.0 | | 99.5 | 98.5 | 98.5 | 97.4 | 22.4 | 22.4 | 22.4 | 23.0 | 22.4 | 23.0 | 21.9 | 23.5 | 23.5 | 23.5 | 6.0 | 64.1 | 63.6 |
| MC3 | 59.5 | 60.5 | 64.6 | 59.5 | 99.5 | 99.5 | | 99.0 | 99.0 | 98.0 | 22.4 | 22.4 | 22.4 | 23.0 | 22.4 | 23.0 | 21.9 | 23.5 | 23.5 | 23.5 | 6.0 | 64.1 | 63.6 |
| MC4 | 59.0 | 60.0 | 64.1 | 59.0 | 98.5 | 98.5 | 99.0 | | 98.0 | 96.9 | 22.4 | 21.9 | 22.4 | 22.4 | 21.9 | 22.4 | 21.9 | 23.5 | 23.5 | 23.5 | 6.0 | 63.6 | 63.1 |
| MC5 | 58.5 | 60.0 | 63.6 | 59.0 | 98.5 | 98.5 | 99.0 | 98.0 | | 96.9 | 21.9 | 21.9 | 21.9 | 22.4 | 21.9 | 22.4 | 21.4 | 23.0 | 23.0 | 23.0 | 6.0 | 63.1 | 62.6 |
| MI17 | 59.5 | 59.5 | 63.6 | 60.0 | 98.0 | 97.4 | 98.0 | 96.9 | 96.9 | | 22.4 | 22.4 | 22.4 | 23.0 | 22.4 | 23.0 | 21.9 | 23.5 | 23.5 | 23.5 | 6.0 | 64.1 | 63.6 |
| MleiPP1 | 25.6 | 23.3 | 25.1 | 26.1 | 22.4 | 22.4 | 22.4 | 22.4 | 21.9 | 22.4 | | 90.3 | 93.2 | 96.6 | 89.9 | 98.5 | 92.3 | 87.4 | 87.0 | 87.0 | 10.0 | 26.5 | 26.5 |
| MleiPP2 | 25.6 | 23.7 | 25.1 | 26.1 | 22.4 | 22.4 | 22.4 | 21.9 | 21.9 | 22.4 | 90.3 | | 97.1 | 91.3 | 99.0 | 90.8 | 96.6 | 89.9 | 89.4 | 89.4 | 9.2 | 25.5 | 25.5 |
| MleiPP3 | 24.6 | 22.4 | 24.1 | 25.1 | 22.4 | 22.4 | 22.4 | 22.4 | 21.9 | 22.4 | 93.2 | 97.1 | | 94.2 | 96.1 | 93.7 | 98.6 | 89.9 | 89.4 | 89.4 | 9.2 | 25.5 | 25.5 |
| MleiPP4 | 25.1 | 22.8 | 24.6 | 25.6 | 23.0 | 23.0 | 23.0 | 22.4 | 22.4 | 23.0 | 96.6 | 91.3 | 94.2 | | 91.8 | 97.6 | 94.2 | 87.0 | 86.5 | 86.5 | 10.0 | 25.5 | 25.5 |
| MleiPP5 | 25.6 | 23.7 | 25.1 | 26.1 | 22.4 | 22.4 | 22.4 | 21.9 | 21.9 | 22.4 | 89.9 | 99.0 | 96.1 | 91.8 | | 90.3 | 97.6 | 89.9 | 89.4 | 89.4 | 9.2 | 25.5 | 25.5 |
| MleiPP6 | 25.1 | 22.8 | 24.6 | 25.6 | 23.0 | 23.0 | 23.0 | 22.4 | 22.4 | 23.0 | 98.5 | 90.8 | 93.7 | 97.6 | 90.3 | | 92.8 | 88.4 | 87.9 | 87.9 | 10.0 | 26.0 | 26.0 |
| MleiPP7 | 24.1 | 22.0 | 23.6 | 24.6 | 21.9 | 21.9 | 21.9 | 21.9 | 21.4 | 21.9 | 92.3 | 96.6 | 98.6 | 94.2 | 97.6 | 92.8 | | 89.4 | 88.9 | 88.9 | 9.2 | 25.0 | 25.0 |
| MleiPP8 | 25.6 | 24.1 | 26.6 | 26.1 | 23.5 | 23.5 | 23.5 | 23.5 | 23.0 | 23.5 | 87.4 | 89.9 | 89.9 | 87.0 | 89.9 | 88.4 | 89.4 | | 99.5 | 99.5 | 8.7 | 26.0 | 26.0 |
| MleiPP9 | 25.6 | 24.1 | 26.6 | 26.1 | 23.5 | 23.5 | 23.5 | 23.5 | 23.0 | 23.5 | 87.0 | 89.4 | 89.4 | 86.5 | 89.4 | 87.9 | 88.9 | 99.5 | | 99.0 | 8.7 | 26.0 | 26.0 |
| MleiPP10 | 25.6 | 24.1 | 26.6 | 26.1 | 23.5 | 23.5 | 23.5 | 23.5 | 23.0 | 23.5 | 87.0 | 89.4 | 89.4 | 86.5 | 89.4 | 87.9 | 88.9 | 99.5 | 99.0 | | 8.7 | 26.0 | 26.0 |
| OBL | 8.3 | 7.6 | 6.9 | 8.3 | 6.0 | 6.0 | 6.0 | 6.0 | 6.0 | 6.0 | 10.0 | 9.2 | 9.2 | 10.0 | 9.2 | 10.0 | 9.2 | 8.7 | 8.7 | 8.7 | | 9.0 | 9.0 |
| OG1 | 77.9 | 74.4 | 75.4 | 76.4 | 64.6 | 64.1 | 64.1 | 63.6 | 63.1 | 64.1 | 26.5 | 25.5 | 25.5 | 25.5 | 25.5 | 26.0 | 25.0 | 26.0 | 26.0 | 26.0 | 9.0 | | 99.5 |
| OG2 | 77.4 | 73.8 | 74.9 | 75.9 | 64.1 | 63.6 | 63.6 | 63.1 | 62.6 | 63.6 | 26.5 | 25.5 | 25.5 | 25.5 | 25.5 | 26.0 | 25.0 | 26.0 | 26.0 | 26.0 | 9.0 | 99.5 | |

[a]Percentage of identity among CaPhs were estimated from the multiple alignment using the MUSCLE tool at the Geneious Prime software (Biomatters).

**Table S2.** Predicted molecular weight (MW) and isoelectric point (pI) values for the *V. parallelum* photoproteins using the software Geneious Prime (Biomatters).

| Velamin | Predicted MW (kDa) | Predicted pI (pH) |
|---|---|---|
| VparPP1 | 24.701 | 4.59 |
| VparPP2 | 24.713 | 4.54 |
| VparPP3 | 24.687 | 4.58 |
| VparPP4 | 24.726 | 4.43 |
| VparPP5 | 24.549 | 4.31 |
| VparPP6 | 24.712 | 4.43 |
| VparPP7 | 24.486 | 4.44 |
| VparPP8 | 24.740 | 4.42 |
| VparPP9 | 24.596 | 4.71 |
| VparPP10 | 24.685 | 4.59 |

**Table S3.** GenBank and PDB accession numbers for CaPhs included in phylogeny estimation and structural comparisons.

| Organism | Photoprotein | Variant | Accession number | PDB |
|---|---|---|---|---|
| *Aequorea victoria* | Aequorin | AEQ1_AEQVI | P07164 | 1SL8 |
| *Aequorea victoria* | Aequorin | AEQ2_AEQVI | P02592 | 1UHK |
| *Aequorea coerulescens* | Aequorin | AEQ_CO | AAO91813 | |
| *Aequorea macrodactyla* | Aequorin | AEQ_MA | AAK02061 | |
| *Aequorea parva* | Aequorin | AEQ_PA | AAK02060 | |
| *Bathocyroe fosteri* | Bathocyroin | BfosPP | (4) | |
| *Beroe abyssicola* | Berovin | Berovin1 | AFE88609 | 4MN0 |
| *Bolinopsis infundibulum* | Bolinopsin | Bolinopsin | CQ975887 | |
| *Bolinopsis infundibulum* | Bolinopsin | Bolinopsin_gr | CS447621 | |
| *Clytia gregaria* | Clytin | Clytin-I | BAG49090 | |
| *Clytia gregaria* | Clytin | Clytin-II | BAG49087 | |
| *Clytia gregaria* | Clytin | Clytin-3 | ADI71937 | 3KPX |
| *Clytia hemisphaerica* | Clytin | Clytin1 | AEP19818 | |
| *Clytia hemisphaerica* | Clytin | Clytin2 | AEP19819 | |
| *Clytia hemisphaerica* | Clytin | Clytin3 | AEP19820 | |
| *Mitrocoma cellularia* | Mitrocomin | MC1 | AIU48026 | |
| *Mitrocoma cellularia* | Mitrocomin | MC2 | AIU48027 | |
| *Mitrocoma cellularia* | Mitrocomin | MC3 | AIU48028 | |
| *Mitrocoma cellularia* | Mitrocomin | MC4 | AIU48029 | |
| *Mitrocoma cellularia* | Mitrocomin | MC5 | AIU48030 | |
| *Mitrocoma cellularia* | Mitrocomin | MC7 | AIU48032 | |
| *Mitrocoma cellularia* | Mitrocomin | MIC17 | AAA29298 | 4NQG |
| *Mnemiopsis leidyi* | Mnemiopsin | MleiPP1 | AFK83778 | 5VP3 |
| *Mnemiopsis leidyi* | Mnemiopsin | MleiPP2 | AFK83779 | |
| *Mnemiopsis leidyi* | Mnemiopsin | MleiPP3 | AFK83780 | |
| *Mnemiopsis leidyi* | Mnemiopsin | MleiPP4 | AFK83781 | |
| *Mnemiopsis leidyi* | Mnemiopsin | MleiPP5 | AFK83782 | |

| | | | | |
|---|---|---|---|---|
| *Mnemiopsis leidyi* | Mnemiopsin | MleiPP6 | AFK83783 | |
| *Mnemiopsis leidyi* | Mnemiopsin | MleiPP7 | AFK83784 | |
| *Mnemiopsis leidyi* | Mnemiopsin | MleiPP8 | AFK83785 | |
| *Mnemiopsis leidyi* | Mnemiopsin | MleiPP9 | AFK83786 | |
| *Mnemiopsis leidyi* | Mnemiopsin | MleiPP10 | AFK83787 | |
| *Obelia geniculata* | Obelin | OG1 | Q8T6Z0 | |
| *Obelia geniculata* | Obelin | OG2 | AIU48034 | |
| *Obelia longuissima* | Obelin | OBL | Q27709 | 1QV1 |
| *Velamen parallelum* | Velamin | VparPP1 | | |
| *Velamen parallelum* | Velamin | VparPP2 | | |
| *Velamen parallelum* | Velamin | VparPP3 | | |
| *Velamen parallelum* | Velamin | VparPP4 | | |
| *Velamen parallelum* | Velamin | VparPP5 | | |
| *Velamen parallelum* | Velamin | VparPP6 | | |
| *Velamen parallelum* | Velamin | VparPP7 | | |
| *Velamen parallelum* | Velamin | VparPP8 | | |
| *Velamen parallelum* | Velamin | VparPP9 | | |
| *Velamen parallelum* | Velamin | VparPP10 | | |

**Table S4.** GenBank accession numbers for the 18S rDNA and COI sequences of Lobates ctenophores included in this study.

| Species | 18S rDNA | COI | Species | 18S rDNA | COI |
|---|---|---|---|---|---|
| *Bathocyroe aff. fosteri* | MW647012 | MW735701 | *Eurhamphaea aff.vexilligera* | MW647038 | MW804236 |
| *Bathocyroe aff. fosteri* | | MW735700 | *Eurhamphaea aff. vexilligera* | MW647039 | MW804237 |
| *Bathocyroe aff. fosteri* | MW647010 | MW735703 | *Eurhamphaea vexilligera* | MW647040 | |
| *Bathocyroe aff. fosteri* | MW647015 | MW735704 | *Eurhamphaea vexilligera* | | MW735765 |
| *Bathocyroe aff. fosteri* | MW647016 | MW735702 | *Eurhamphaea vexilligera* | MW647042 | MW735768 |
| *Bathocyroe aff. longigula* | | MW735707 | *Kiyohimea aurita* | MW647043 | |
| *Bathocyroe aff. longigula* | MW647009 | MW735705 | *Kiyohimea usagi* | MW647044 | MW735782 |
| *Bathocyroe aff. longigula* | | MW735706 | *Kiyohimea usagi* | MW647045 | MW735783 |
| *Bathocyroe aff. fosteri* | MW647008 | | *Lampocteis cruentiventer* | MW647046 | |
| *Bathocyroe fosteri* | MW647011 | | *Lampocteis cruentiventer* | MW647047 | MW735788 |
| *Bathocyroe fosteri* | MW647013 | | *Lampocteis cruentiventer* | MW647048 | MW735791 |
| *Bathocyroe fosteri* | MW647014 | | *Lampocteis cruentiventer* | | MW735789 |
| *Bolinopsis aff. infundibulum* | MW647017 | | *Lampocteis cruentiventer* | MW647049 | MW735790 |
| *Bolinopsis aff. infundibulum* | MW647018 | | *Lampocteis sp.* | MW647050 | MW735792 |
| *Bolinopsis microptera* | | MW735734 | *Lampocteis sp.* | MW647051 | MW735793 |
| *Bolinopsis microptera* | | MW735735 | *Lampocteis sp.* | MW647052 | |
| *Bolinopsis aff. infundibulum* | MW647020 | | *Lampocteis sp.* | | MW735796 |
| *Bolinopsis microptera* | MW647021 | MW735733 | *Lampocteis sp.* | | MW735794 |
| *Bolinopsis aff. infundibulum* | MW647022 | | *Lampocteis sp.* | MW647053 | MW735795 |
| *Bolinopsis aff. infundibulum* | MW647023 | | *Leucothea pulchra* | MW647054 | MW735797 |
| *Bolinopsis aff. infundibulum* | MW647024 | | *Leucothea pulchra* | MW647055 | MW735798 |
| *Bolinopsis microptera* | | MW735736 | *Leucothea pulchra* | MW647056 | |
| *Bolinopsis microptera* | | MW735737 | *Leucothea pulchra* | AF293688 | MW735799 |
| *Bolinopsis microptera* | MW647019 | MW735732 | *Mnemiopsis leidyi* | MW647058 | MW735801 |
| *Bolinopsis aff. vitrea* | MW647025 | | *Mnemiopsis leidyi* | | MW735802 |
| *Bolinopsis aff. vitrea* | MW647027 | | *Mnemiopsis leidyi* | MW647059 | |
| *Bolinopsis aff. vitrea* | | MW735739 | *Ocyropsis aff.crystallina* | | MW735803 |
| *Bolinopsis aff. vitrea* | MW647026 | | *Ocyropsis aff.crystallina* | | MW735804 |
| *Bolinopsis aff. vitrea* | | MW735738 | *Ocyropsis aff.crystallina* | | MW735805 |
| *Bolinopsis ashleyi* | | MW735740 | *Ocyropsis aff.crystallina* | | MW735806 |
| *Bolinopsis infundibulum* | AF293687 | MW735741 | *Ocyropsis aff.crystallina* | | MW735816 |
| *Bolinopsis infundibulum* | MW647029 | | *Ocyropsis aff.crystallina* | | MW735817 |
| *Bolinopsis infundibulum* | | MW735742 | *Ocyropsis aff.crystallina* | | MW735818 |
| *Bolinopsis mikado* | | MW735743 | *Ocyropsis aff. crystallina* | MW647060 | |
| *Bolinopsis mikado* | | MW735744 | *Ocyropsis crystallina crystallina* | AF293690 | MW735807 |
| *Bolinopsis vitrea* | | MW735745 | *Ocyropsis crystallina guttata* | AF293691 | MW735808 |
| *Bolinopsis vitrea* | MW647028 | | *Ocyropsis maculata maculata* | MW647064 | |
| *Cestum aff. veneris* | | MW735751 | *Ocyropsis maculata maculata* | MW647065 | MW735810 |
| *Cestum veneris* | AF293692 | | *Ocyropsis maculata maculata* | MW647066 | MW735811 |
| *Cestum aff. veneris* | MW647032 | | *Ocyropsis maculata maculata* | | MW735812 |
| *Cestum aff. veneris* | MW647033 | | *Ocyropsis maculata maculata* | MW647067 | MW735813 |
| *Cestum aff. veneris* | | MW735749 | *Ocyropsis maculata maculata* | MW647061 | MW735809 |
| *Cestum aff. veneris* | MW647034 | MW735752 | *Ocyropsis maculata maculata* | | MW735815 |
| *Cestum aff. veneris* | MW647035 | MW735750 | *Ocyropsis maculata maculata* | AF293689 | MW735814 |
| *Deiopea aff. kaloktenota* | MW647037 | | *Pleurobrachia bachei* | | MW735819 |
| *Deiopea sp* | | MW735755 | *Thalassocalyce inconstans* | MW647070 | MW735825 |
| *Deiopea sp* | | MW735756 | *Thalassocalyce inconstans* | MW647071 | MW735826 |
| *Deiopea aff. kaloktenota* | | MW735757 | *Thalassocalyce inconstans* | MW647072 | MW735827 |
| *Deiopea aff. kaloktenota* | | MW735758 | *Thalassocalyce inconstans* | MW647073 | MW735828 |
| *Deiopea aff. kaloktenota* | MW647036 | MW735761 | *Velamen parallelum* | | MW735832 |
| *Deiopea aff. kaloktenota* | | MW735760 | *Velamen parallelum* | AF293693 | MW735830 |
| *Deiopea aff. kaloktenota* | | MW735759 | *Velamen parallelum* | MW647074 | MW735831 |
| *Eurhamphaea vexilligera* | MW647041 | MW735766 | *Velamen parallelum (this study)* | OQ463859 | PP490733 |

**Table S5.** Raw data from light emission assays of velamins in different pH (Fig. 3A).

| | | Integral (× $10^{-6}$ counts)[a] | | |
|---|---|---|---|---|
| **Buffer[b]** | **pH** | **α-velamin** | **β-velamin** | **γ-velamin** |
| NaPi | 6.0 | 0.040 ± 0.004 | 0.060 ± 0.002 | 0.010 ± 0.002 |
| | 6.5 | 0.020 ± 0.001 | 0.020 ± 0.002 | 0.080 ± 0.001 |
| | 7.0 | 0.17 ± 0.03 | 0.120 ± 0.002 | 0.14 ± 0.01 |
| | 7.5 | 0.050 ± 0.003 | 0.22 ± 0.02 | 0.33 ± 0.03 |
| TRIS | 7.5 | 0.90 ± 0.05 | 0.75 ± 0.09 | 0.44 ± 0.03 |
| | 8.0 | 2.0 ± 0.3 | 1.4 ± 0.2 | 0.8 ± 0.2 |
| | 8.5 | 3.5 ± 0.4 | 1.6 ± 0.2 | 0.94 ± 0.03 |
| | 9.0 | 2.8 ± 0.1 | 1.7 ± 0.3 | 0.83 ± 0.02 |
| Gly | 9.0 | 2.2 ± 0.2 | 1.6 ± 0.1 | 1.0 ± 0.2 |
| | 9.5 | 1.5 ± 0.1 | 0.81 ± 0.03 | 0.59 ± 0.02 |
| | 10 | 0.44 ± 0.04 | 0.25 ± 0.02 | 0.190 ± 0.004 |

[a]Velamins were regenerated in 50 mM Tris buffer pH 9 by adding 0.5 µM purified apovelamin, 1 µM coelenterazine (NanoLight) and 10 mM EDTA into 1.5 mL amber microtubes and kept at 4ºC for 12h. For light emission assays, 500 µL of 50 mM buffer supplemented with 0.5 mM $CaCl_2$ was added to 1 µL of velamins and light emission was recorded during 10s using the luminometer Sirius L (Berthold). Integral values and standard deviation were estimated from technical replicates. [b]Different buffers were used in chemiluminescent assays, including sodium phosphate (pH range 6-7.5), Tris (pH range 7.5-9), and glycine (pH range 9-10).

**Table S6.** Raw data from light emission assays to assess the effects of different incubation times during the regeneration of velamins on the light emission (Fig. 3C).

| Incubation Time (h) | Integral (× $10^{-6}$ counts)[a] | | |
|---|---|---|---|
| | α-velamin | β-velamin | γ-velamin |
| 0 | 2.0 ± 0.2 | 1.7 ± 0.2 | 1.7 ± 0.1 |
| 2 | 2.3 ± 0.1 | 2.0 ± 0.1 | 1.1 ± 0.1 |
| 4 | 4.0 ± 0.1 | 3.4 ± 0.1 | 1.7 ± 0.1 |
| 8 | 5.5 ± 0.2 | 4.2 ± 0.1 | 2.4 ± 0.1 |
| 10 | 5.6 ± 0.3 | 4.4 ± 0.4 | 2.5 ± 0.1 |
| 12 | 6.0 ± 0.2 | 4.3 ± 0.1 | 2.6 ± 0.1 |
| 16 | 5.3 ± 0.1 | 4.1 ± 0.2 | 2.0 ± 0.1 |
| 22 | 5.3 ± 0.4 | 4.1 ± 0.3 | 2.3 ± 0.1 |
| 24 | 5.1 ± 0.4 | 4.0 ± 0.1 | 2.3 ± 0.1 |

[a]Velamins were regenerated in 50 mM Tris buffer pH 9 by adding 0.5 μM purified apovelamin, 1 μM coelenterazine (NanoLight) and 10 mM EDTA into 1.5 mL amber microtubes and kept at 4ºC during different times of incubation (0, 2, 4, 8, 10, 12, 16, 22 and 24h). Light emission assays were performed by adding 500 μL of 50 mM Tris buffer pH 9 supplemented with 0.5 mM $CaCl_2$ to 1 μL of velamins. Light emission was recorded during 10s using the luminometer Sirius L (Berthold). Integral values and standard deviation were estimated from technical replicates.

**Table S7.** Raw data from light emission assays to assess the response of α-, β-, and γ-velamins to calcium (Fig. 3D).

| $[Ca^{2+}]$ (M) | Integral (× $10^{-6}$ counts)[a] | | |
|---|---|---|---|
| | **α-velamin** | **β-velamin** | **γ-velamin** |
| $10^{-8}$ | 0.018 ± 0.008 | 0.033 ± 0.005 | 0.06 ± 0.02 |
| $10^{-7}$ | 0.012 ± 0.006 | 0.030 ± 0.007 | 0.027 ± 0.001 |
| $10^{-6}$ | 0.007 ± 0.002 | 0.061 ± 0.007 | 0.04 ± 0.01 |
| $10^{-5}$ | 2.9 ± 0.1 | 3.1 ± 0.3 | 2.0 ± 0.1 |
| $10^{-4}$ | 3.02 ± 0.04 | 3.4 ± 0.2 | 1.9 ± 0.1 |
| $10^{-3}$ | 2.8 ± 0.1 | 2.7 ± 0.1 | 1.7 ± 0.1 |
| $10^{-2}$ | 2.15 ± 0.03 | 2.0 ± 0.4 | 1.25 ± 0.04 |

[a]Velamins were regenerated in 50 mM Tris buffer pH 9 by adding 0.5 µM purified apovelamin, 1 µM coelenterazine (NanoLight) and 10 mM EDTA into 1.5 mL amber microtubes and kept at 4ºC for 12h. Light emission assays were performed by adding 500 µL of 50 mM Tris buffer pH 9 supplemented with $CaCl_2$ (concentrations from $10^{-2}$ to $10^{-7}$ M) to 1 µL of velamins. Light emission was recorded during 10s using the luminometer Sirius L (Berthold). Integral values and standard deviation were estimated from technical replicates.

**Table S8.** Raw data from light emission assays to assess the thermostability of apovelamins (Fig. 4).

| Incubation Time (min) | α-velamin[a] | | | |
|---|---|---|---|---|
| Normalized Integral (%)/Temperature (ºC) | | | | |
| 0[b] | 100 ± 3/0ºC | | | |
| 10 | 73 ± 2/25ºC | 76 ± 4/37ºC | 71 ± 6/50ºC | 21 ± 1/75ºC |
| 20 | 86 ± 7/25ºC | 68 ± 5/37ºC | 68 ± 4/50ºC | 16 ± 1/75ºC |
| 30 | 88 ± 4/25ºC | 76 ± 3/37ºC | 55 ± 2/50ºC | 11 ± 1/75ºC |
| 60 | 81 ± 2/25ºC | 81 ± 1/37ºC | 34 ± 1/50ºC | 6 ± 1/75ºC |
| β-velamin | | | | |
| 0 | 48 ± 1/0ºC | | | |
| 10 | 70 ± 3/25ºC | 55 ± 2/37ºC | 55 ± 1/50ºC | 18 ± 1/75ºC |
| 20 | 54 ± 2/25ºC | 67 ± 2/37ºC | 33 ± 3/50ºC | 11 ± 1/75ºC |
| 30 | 61 ± 2/25ºC | 64 ± 3/37ºC | 39 ± 3/50ºC | 8 ± 1/75ºC |
| 60 | 61 ± 1/25ºC | 54 ± 6/37ºC | 28 ± 1/50ºC | 6 ± 1/75ºC |
| γ-velamin | | | | |
| 0 | 42 ± 3/0ºC | | | |
| 10 | 47 ± 2/25ºC | 51 ± 2/37ºC | 30 ± 1/50ºC | 10 ± 1/75ºC |
| 20 | 38 ± 2/25ºC | 51 ± 2/37ºC | 30 ± 1/50ºC | 7 ± 1/75ºC |
| 30 | 55 ± 1/25ºC | 41 ± 2/37ºC | 28 ± 1/50ºC | 3 ± 1/75ºC |
| 60 | 51 ± 4/25ºC | 44 ± 2/37ºC | 16 ± 1/50ºC | 2 ± 1/75ºC |

[a]Apovelamins were incubated at 25, 37, 50, and 75°C for different periods of time (0, 10, 20, 30, and 60 min) in 50 mM Tris buffer pH 9 containing 10 mM EDTA. Then, samples were placed on ice for 30 min and regenerated in the presence of 1 µM coelenterazine (NanoLight) into 1.5 mL amber microtubes kept at 4ºC for 12h. Light emission assays were performed by adding 500 µL of 50 mM Tris buffer pH 9 supplemented with 0.5 mM $CaCl_2$ to 1 µL of velamins. Light emission was recorded during 10s using the luminometer Sirius L (Berthold). Integral values and standard deviation were estimated from technical replicates. [b]All values were normalized using α-cestidin as standard: chemiluminescence integral (3.3 ± 0.4) × $10^6$ counts, determined from a sample regenerated at 0ºC and at 0 min.

**Table S9.** Raw data from light emission assays to assess the thermal stability of α-velamin at 37ºC (Fig. S5A).

| **Exposition Time (min)** | **Integral (× $10^{-5}$ counts)[a]** |
|---|---|
| 1 | 3.2 ± 0.2 |
| 5 | 1.5 ± 0.08 |
| 10 | 0.08 ± 0.04 |
| 15 | 0.008 ± 0.005 |

[a]α-velamin was regenerated in 50 mM Tris buffer pH 9 containing 10 mM EDTA and 1 μM coelenterazine (NanoLight) into 1.5 mL amber microtubes, kept at 4ºC for 12h. Then, α-velamin samples were incubated at 37ºC during 1, 5, 10, and 15 min, followed by light emission assays performed by adding 500 μL of 50 mM Tris buffer pH 9 supplemented with 0.5 mM $CaCl_2$ to 1 μL of velamins. Light emission was recorded during 10s using the luminometer Sirius L (Berthold). Integral values and standard deviation were estimated from technical replicates.

**Table S10.** Raw data from light emission assays to assess the photoinactivation of α-velamin by the visible light (Fig. S5B).

| Exposition Time (min) | Integral (× $10^{-5}$ counts)[a] |
|---|---|
| 0 | 6.9 ± 0.1 |
| 5 | 5.3 ± 0.8 |
| 10 | 3.4 ± 0.5 |
| 25 | 1.4 ± 0.05 |
| 60 | 0.3 ± 0.04 |

[a]α-velamin was regenerated in 50 mM Tris buffer pH 9 containing 10 mM EDTA and 1 µM coelenterazine (NanoLight) into 1.5 mL amber microtubes, kept at 4ºC for 12h. Then, α-velamin samples were exposed to visible light during 5, 10, 25, and 60 min, followed by light emission assays performed by adding 500 µL of 50 mM Tris buffer pH 9 supplemented with 0.5 mM $CaCl_2$ to 1 µL of velamins. Light emission was recorded during 10s using the luminometer Sirius L (Berthold). Integral values and standard deviation were estimated from technical replicates.

**Movie S1 (separate file).** *Velamen parallelum* specimens observed close to the Alcatrazes Archipelago (São Paulo state, Brazil).

**Movie S2 (separate file).** Chemiluminescence reaction of α-velamin.

**Movie S3 (separate file).** Structural comparison at the N-termini of CaPhs from ctenophores and hydromedusae.

**Movie S4 (separate file).** Superimposed structures from photoproteins highlighting the replacement of the conserved triad His-Trp-Tyr, found in hydromedusan CaPhs, with the equivalent Phe-Trp-Tyr in ctenophore CaPhs.

**Movie S5 (separate file).** Conserved triad Arg40-Asp157-Tyr203 (VparPP2) proposed to be involved in light emission in ctenophore CaPhs.
